# Deposit-feeding worms control subsurface ecosystem functioning in intertidal sediment with strong physical forcing

**DOI:** 10.1101/2022.04.06.487375

**Authors:** Longhui Deng, Christof Meile, Annika Fiskal, Damian Bölsterli, Xingguo Han, Niroshan Gajendra, Nathalie Dubois, Stefano M. Bernasconi, Mark A. Lever

**Author notes:** **Correspondence:** Longhui Deng, Mark Alexander Lever. Department of Microbial Ecology, German Federal Institute of Hydrology (BfG), Am Mainzer Tor 1, 56068, Koblenz, Germany (Annika Fiskal). Forest Soils and Biogeochemistry, Swiss Federal Institute for Forest, Snow and Landscape Research WSL, 8903 Birmensdorf, Switzerland. **Author Contributions:** L.D. and M.A.L. designed research; L.D., A.F., D.B., and M.A.L. conducted field investigations; L.D., C.M., A.F., D.B., X.H., N.G., N.B., S.M.B., and M.A.L. performed measurements and data analyses; C.M. and M.A.L. supervised the project; L.D. and M.A.L. wrote the manuscript with inputs from all co-authors. **Competing Interest Statement:** Authors declare that they have no competing interests.

## Abstract

Intertidal sands are global hotspots of terrestrial and marine carbon cycling with strong hydrodynamic forcing by waves and tides and high macrofaunal activity. Yet, the relative importance of hydrodynamics and macrofauna in controlling these ecosystems remains unclear. Here we compare bacterial, archaeal, and eukaryotic communities in upper intertidal sands dominated by subsurface deposit-feeding worms (*Abarenicola pacifica*) to adjacent worm-free areas. We show that hydrodynamic forcing controls organismal assemblages in surface sediments, while in deeper layers selective feeding by worms on fine, algae-rich particles strongly decreases the abundance and richness of all three domains. In these deeper layers, bacterial and eukaryotic network connectivity decreases, while percentages of taxa involved in degradation of refractory organic macrostructures, oxidative nitrogen and sulfur cycling, and macrofaunal symbioses, increase. Our findings reveal macrofaunal activity as the key driver of ecosystem functioning and carbon cycling in intertidal sands below the mainly physically controlled surface layer.

**Significance Statement:** Hydrodynamics and bioturbation are the main forces controlling chemical exchanges between sediment and seawater in coastal environments. However, little is known about the relative impact of both processes on sediment biological communities. We show that intertidal sand ecosystems dominated by lugworms can be divided into vertically distinct hydrodynamically and biologically controlled layers. Hydrodynamic forcing controls biological communities in surface layers by regulating organic carbon and electron acceptor inputs. By contrast, lugworms structure subsurface ecosystems through the selective consumption of fine particles, which diminishes microbial and eukaryotic populations and weakens ecological networks, while promoting the burial of, mostly terrestrial, macrodetritus. Our study demonstrates that globally distributed marine invertebrates control intertidal sand ecosystems below the physically controlled surface layer.

## Introduction

Burrowing macrofauna have impacted seafloor habitats and altered the marine oxygen and sulfur cycles since their first appearance during the Proterozoic-Cambrian transition (1). Today, these organisms are ubiquitous in seafloor sediments except under conditions of oxygen-depleted bottom water, toxic hydrogen sulfide concentrations, or high temperatures (2). These macrofauna influence sediment matrices through a process called “bioturbation”, which includes the processes of ‘ventilation’ and ‘reworking’ (3). During ventilation fauna flush their burrows with oxygenated seawater to breathe or feed on suspended particles, promoting water exchanges between burrows and surrounding sediment (‘bioirrigation’) (4). Reworking refers to the movement of sediment particles as a result of macrofaunal locomotion, burrowing, or feeding (5).

Ventilation and reworking influence microbially mediated processes by modulating the redox state and gradients of electron acceptors and donors within sediment. By introducing the high-energy electron acceptors oxygen and nitrate from overlying water and creating redox oscillations, ventilation stimulates the microbial oxidation of organic carbon (OC) (4), coupled nitrification–denitrification reactions (6), and oxidative removal of potentially inhibitory microbial end products, such as reduced metals, sulfide, and ammonium (7). In contrast, reworking can fuel microbial activities in deeper layers by introducing freshly deposited organic matter (8), nutrient-rich secretes, e.g. mucopolysaccharides (9), and metal oxides (10), and represents an important control of OC mineralization and burial in marine sediment (1-4, 11, 12).

Comparatively less is known about the interactions among sediment macro-, meio-, and microbiota, even though these interactions are crucial to understanding the flow of carbon through benthic food webs (12). Macrofauna alter the abundance, diversity, and community structure of micro- and meiobiota locally in burrow walls, feeding pockets, and feces (13, 14). More recent studies on subtidal sediments show that the influence of macrofaunal activity is not restricted to burrows but controls microbial community structure throughout the entire bioturbated layer (15, 16). This results in highly similar microbial communities across bioturbated coastal and continental shelf sediments that differ in macrofaunal species compositions, sedimentation rate, sediment geochemistry, and lithology (16). Hereby the dominant lineages in bioturbated surface sediment differ from those that dominate non-bioturbated surface or subsurface marine sediment.

Compared to subtidal habitats, intertidal sediments experience stronger daily fluctuations in tidal and wave action, temperature, and water content (17). Despite these frequent perturbations, intertidal sediments have among the highest benthic microalgal productivities (10^3^- 10^4^ mmol C m^2^ y^-1^) and OC decomposition rates (10^3^-10^5^ mmol C m^2^ y^−1^) across seafloor ecosystems (18-20), and frequently host high macrofaunal biomass (21). Both hydrodynamic forcing and bioturbation impact O_2_ dynamics, OC remineralization, and nutrient fluxes in intertidal sediments (20, 22). In addition, bioturbation changes food-web structures by promoting microbial grazing (13), symbioses (23) and meiofauna-microorganism interactions (24). Despite the known impacts of hydrodynamic forcing and bioturbation, it remains largely unknown how both forces compare, or interact with each other, in intertidal ecosystems.

Here we explore the impact of physical forcing and bioturbation on intertidal carbon cycling and community functioning based on sandy sediment of False Bay, Washington, USA. Study sites were dominated by lugworms, a group of polychaetes with worldwide distributions in coastal sediments (25) that account for up to 30% of benthic biomass in intertidal sediments (21). Past research on *Abarenicola pacifica*, the dominant lugworm species in False Bay, led to the now broadly recognized concept of “microbial gardening”, which originally described the stimulation of microbial growth by *A. pacifica* in subsurface sediments (13).

We investigate the relative importance of hydrodynamic forcing and macrofaunal activity based on adjacent sediment plots with and without lugworms that are subject to similar hydrodynamic forcing and geochemical conditions. To this end, we integrate analyses of sedimentary features (grain size, biogenic structures), porewater geochemical gradients (electron acceptors, metabolic end products, redox potential), OC (content, isotopic compositions, carbon-to-nitrogen ratios (C:N), chemical reactivity), and bacterial, archaeal, and eukaryotic DNA sequences (abundance, richness, sulfur (S)- and nitrogen (N)-cycling genes, community structure) with modeled rates of hydrodynamics and macrofauna-driven porewater and particle mixing. Based on our findings, we propose that lugworms, and possibly other burrowing macrofauna, drive intertidal sand ecosystems below the physically controlled surface layer.

## Results

### Impact of A. pacifica on sediment particle distributions

Previous studies indicate that *A. pacifica* forms “J-shaped” burrows in sandy sediments (Fig. 1A), at the end of which it selectively ingests fine particles and thereby causes a constant downward movement of sediment (13, 25). The digested fine particles are expelled back to the sediment surface and homogenized with surrounding sediment during tidal inundation. Matching this feeding behavior, we observed on average coarser grain sizes in *A. pacifica*-inhabited sediments compared to worm-free controls (*p*<0.05, pairwise t test), with maximum grain sizes at ∼15-25 cm (Fig. 1A). A further consequence of particle preference and feeding-related downward movement of sediment by *A. pacifica* was that sediments below burrows (∼20-25 cm) were highly enriched in woody debris, and to a lesser degree macroalgae <u>(Ulvophyceae)</u> detritus and bivalve shells. These biogenic macrostructures were largely absent in control sediments. Based on observed living depths of *A. pacifica* and distributions of macrostructures, measured sediment grain size distributions, and modelled coefficients of physical and biological mixing (details see later), we divide sediments with worms into **(1)** physically and biologically impacted layers (**PBL**; 0-12.5 cm) that overlie the main living depth of *A. pacifica*; **(2)** biologically impacted layers (**BL**; 12.5-25 cm), where lugworms perform most of their feeding and reworking; horizontal burrows, which correspond to the main living depths of *A. pacifica*, were typically located in the upper half of the BL; and **(3)** undisturbed layers (**UL**; >25 cm). Controls without lugworms were divided into physically impacted layers (**PL**; 0-12.5 cm), where significant sediment mixing and porewater exchanges due to tide-related water movements were detectable, and undisturbed layers below (**UL**; >12.5 cm).

**Fig. 1.**
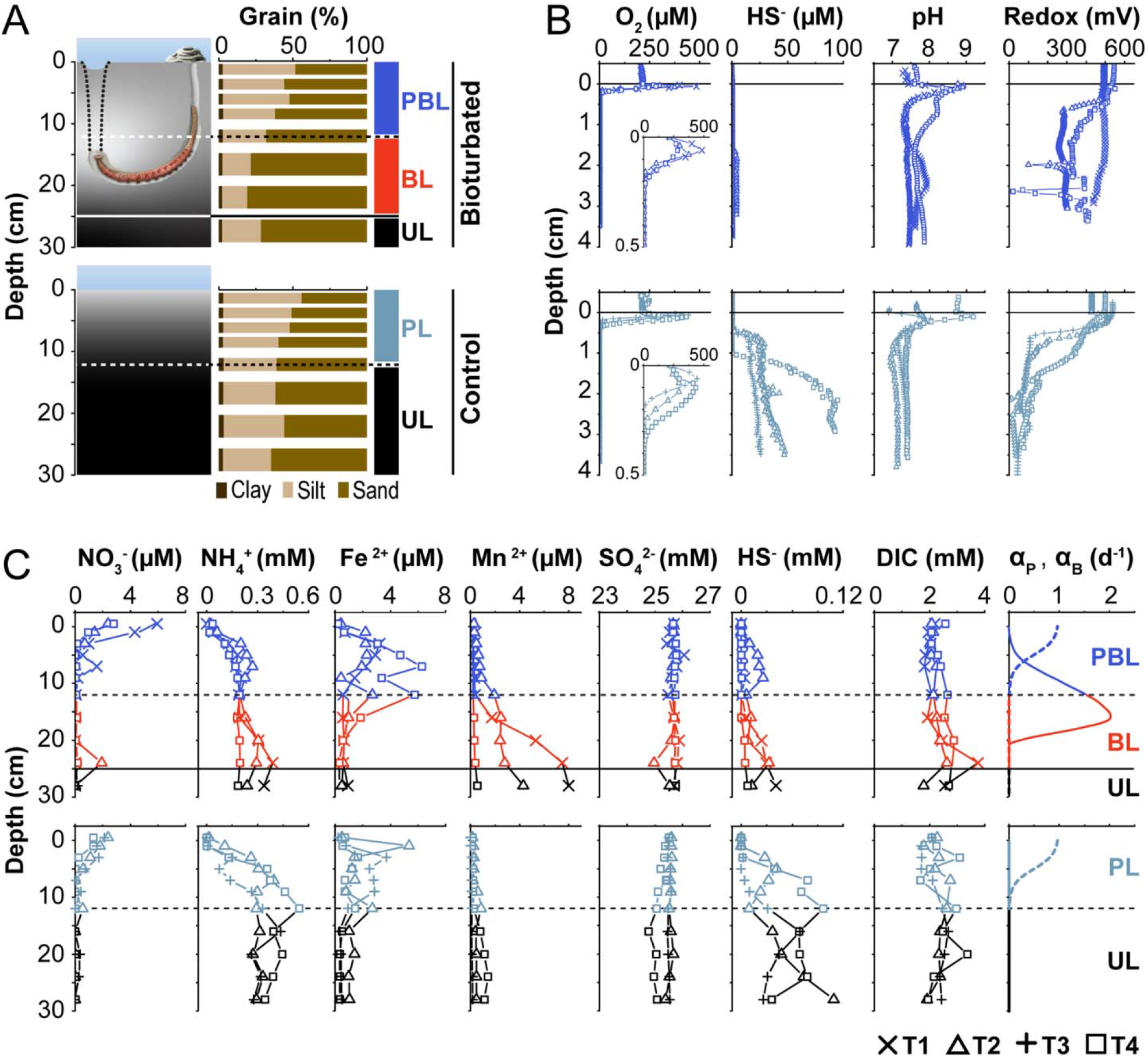
Gradients of porewater geochemistry and porewater exchanges in *Abarenicola pacifica*-inhabited (bioturbated, upper row) and control sediment (lower row). (A) Schematic diagrams of *A. pacifica*-inhabited and control sediments, with measured grain size distributions and proposed sediment zonation of physical and biological mixing. Physically impacted layers (PL), physically and biologically impacted layers (PBL), biologically impacted layers (BL), and undisturbed layers (UL) were determined based on modelled coefficients of physical and biological mixing and observed *A. pacifica* living depths (see text for details). The dotted lines denote the feeding funnel of *A. pacifica*. Fecal piles at the burrow opening attest to the high rates at which *A. pacifica* transports ingested subsurface sediments to the sediment surface. (B) Microsensor profiles of O_2_, HS^-^, pH, and redox potential. (C) Porewater profiles of NO_3_^-^, SO_4_^2-^, HS^-^, NH_4_^+^, DIC, Fe^2+^, and Mn^2+^, and modeled coefficients of physical porewater exchanges (αP, dashed line) and bioirrigation (αB, solid line). Symbol codes: T1 (**×**), T2 (△), T3 (+), and T4 (□) denote data from different time points. Molar concentrations in (B) and (C) are per volume of sediment porewater.

### Impact of lugworm bioturbation on porewater geochemical gradients

Microsensor profiles reveal shallower oxygen (O_2_) penetration, lower hydrogen sulfide (HS^-^) accumulation, and more variable but higher pH and redox potentials in bioturbated compared to control treatments (Fig. 1B). While dissolved O_2_ concentration peaks near the sediment-water interface likely reflect microphytobenthic photosynthesis, O_2_ penetrations are shallow (<0.5 cm) in both treatments. HS^-^ concentrations in the top 4 cm remain low (<5 µM) in bioturbated sediments but increase to 20-100 µM in controls. pH values decrease with depth and are slightly higher at >1 cm in bioturbated (7.6±0.2) than in control sediments (7.3±0.1). Redox potentials decrease from 400-600 mV in overlying water to 333±84 mV and 86±56 mV at >1 cm in bioturbated and control sediments, respectively, indicating more oxidizing conditions in *A. pacifica* treatments.

Bioturbated sediments generally have higher concentrations of electron acceptors (nitrate (NO_3_^-^), sulfate (SO_4_^2-^)) and end products of microbial metal cycling (Fe^2+^, Mn^2+^), but lower accumulations of other reduced end products (ammonium (NH_4_^+^), hydrogen sulfide (HS^-^)) compared to controls (Fig. 1C). NO_3_^-^ concentrations are higher in surface sediments of worm treatments (∼3-8 µM) compared to controls (1-3 µM), and typically decrease to ∼0 µM at >5 cm in both treatments. SO_4_^2-^ concentrations show minor depth variations but are on average slightly higher in bioturbated sediments (25.7±0.2 vs. 25.4±0.2 mM). NH_4_^+^ and HS^-^ concentrations show bimodal profiles with subsurface minima at ∼12-16 cm, and are significantly lower in bioturbated sediments (both *p*<0.05, hereafter all *p* values are based on Welch’s t test unless stated otherwise). Fe^2+^ concentrations remain in the low micromolar range but are slightly higher in the top 0-15 cm of bioturbated compared to control sediments, and are low in >15 cm of both treatments. Mn^2+^ concentrations also remain in the low micromolar range, but increase in deeper layers, especially in the BL. DIC concentrations fluctuate between ∼1-3 mM in both treatments.

Porewater exchange intensities (rightmost panel in Fig. 1C) are estimated from model simulations matching the DIC concentration profiles in manipulation experiments with and without worms (see Methods), and indicate physically induced porewater exchanges (α_P_) in the top ∼10 cm of sediment. Biologically induced porewater mixing (α_B_) indicates active pumping of seawater into worm burrow that causes bioirrigation to ∼10-20 cm in bioturbated treatments, but not in controls.

### Impact of reworking on distributions of OC pools and sources

Depth profiles of OC quantity (total organic carbon (TOC)), OC source (C:N, δ^13^C-TOC), and OC chemical reactivity (chlorophyll *a*, pheopigments, freshness index) indicate that *A. pacifica* modifies the distributions of OC fractions (Fig. 2). This is confirmed by copy numbers of ribulose-1,5-bisphosphate carboxylase genes (*rbcL*) of *Ochrophyta* (mainly diatoms based on 18S rRNA gene sequencing) and vascular plants. TOC and C:N in *A. pacifica* treatments are constantly low in the PBL (0.13±0.04% and 8±2, respectively), increase to ∼0.6% and ∼30 at the bottom of the BL, and decrease to ∼0.1-0.2 % and ∼10-20 below in the UL, respectively (Fig. 2A). δ^13^C-TOC generally decreases from ∼ -19‰ at 1 cm to ∼ -27‰ at the bottom of the BL, with a slight local increase (∼ -21‰) at ∼16 cm. The maxima in TOC and C:N and minima in δ^13^C-TOC at the bottom of the BL correspond to high-sand layers below *A. pacifica* burrows. In comparison, TOC, C:N, and δ^13^C-TOC in controls are more variable and show no consistent trends related to depth.

**Fig. 2.**
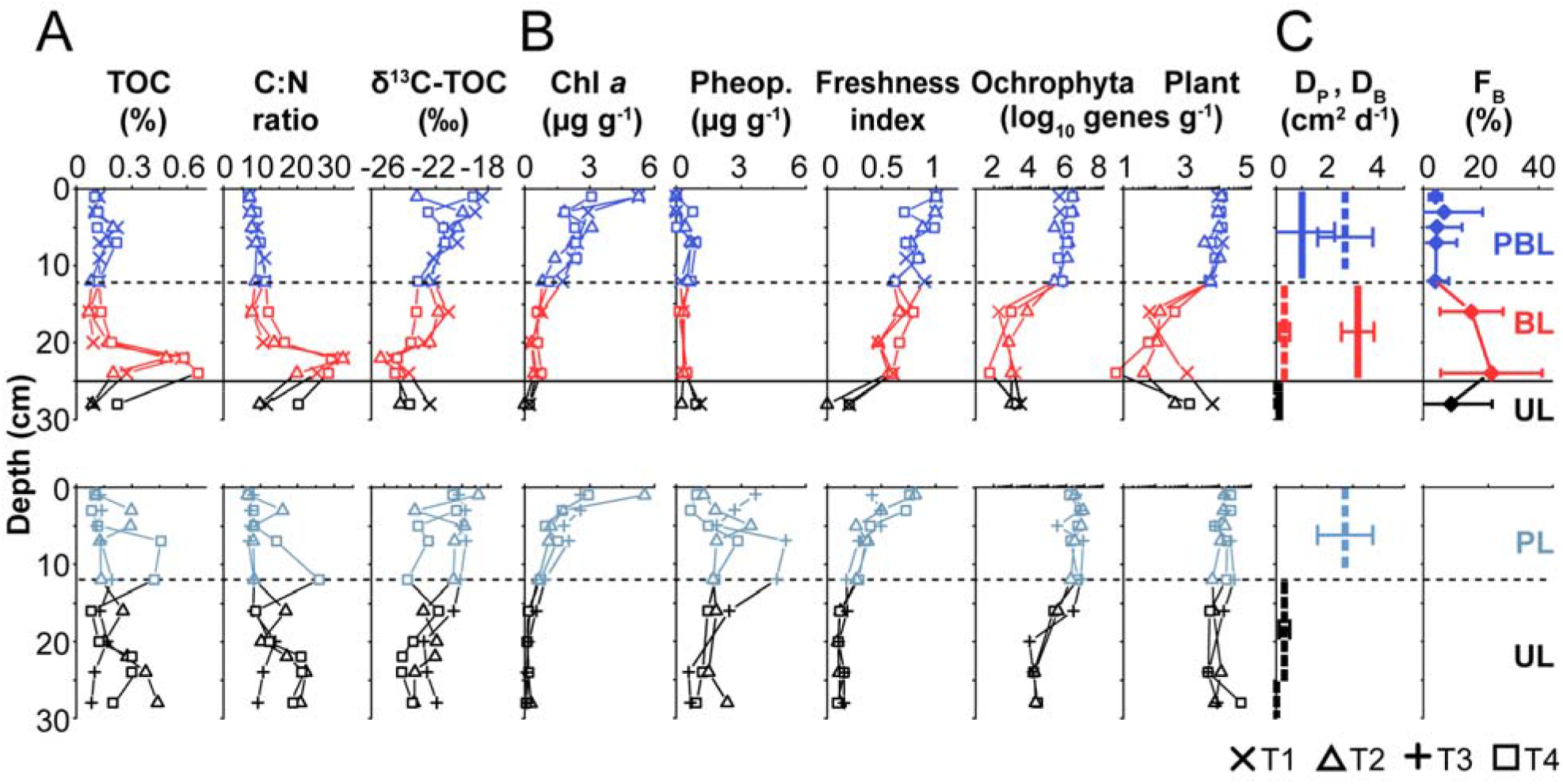
Distributions, sources, and reactivities of OC and particle mixings in bioturbated (top row) and control (bottom row) sediments. (A) Bulk OC quantity (TOC (% dry weight)) and source indicators (C:N, δ^13^C-TOC). (B) Indicators of reactive OC pools: chlorophyll *a* (chl *a*) and pheopigments (pheop.; both g^-1^ wet sediment), freshness index, and gene abundances (*rbc*L) of *Ochrophyta* and vascular plants (as log10 *rbc*L copies g^-1^ wet sediment). (C) Modelled coefficients of physical (D_P_, dashed line) and biological (D_B_, solid line) particle mixing and feeding intensity (F_B_).

Distributions of labile OC fractions also differ clearly between treatments (Fig. 2B). Chl *a* values decrease with depth, with slightly slower depth attenuation and consequently higher chl *a* at 12.5-25 cm in bioturbated treatments. By contrast, pheopigments (chl *a* degradation products) are markedly lower in bioturbated sediments compared to controls (0.5±0.7 vs. 2.0 ±1.2 µM, *p*<0.001). The freshness index (ratio of chl *a* to total pigments) decreases more gradually and is higher (*p*<0.001) in the top 25 cm of bioturbated sediment compared to controls. *rbc*L abundances of *Ochrophyta* and vascular plants in bioturbated sediments are stable around 10^6^ and 10^4^ copies g^-1^ from 0-12.5 cm, respectively, and drop to ≤ 10^2^ genes g^-1^ in the BL. *rbc*L abundances of *Ochrophyta* remain low, whereas those of vascular plants increase again by 1-2 orders of magnitude in the bottom of the BL or UL. In controls, *Ochrophyta* and vascular-plant genes are slightly higher (∼10^6^-10^7^ and ∼10^4^-10^5^ genes g^-1^, respectively) from 0-12.5 cm and decrease less with depth, remaining at *∼*10^4^ and ∼10^4^-10^5^ genes g^-1^, respectively.

Coefficients of physically (D_P_) and biologically (D_B_) induced particle mixing (see Methods for details) suggest a strong decrease in D_P_ from surface to subsurface sediments, and a clearly increased importance of D_B_ in subsurface layers with *A. pacifica* (Fig. 2C). Matching the latter, feeding intensity (F_B_; see Methods) increases significantly in the BL.

### Influence of lugworm bioturbation on gene abundances

Abundance profiles of bacterial and archaeal 16S rRNA and eukaryotic 18S rRNA genes, and of functional genes involved in microbial S- and N-cycling (*dsr*B: *ß* subunit of dissimilatory sulfite reductase; *sox*B: *ß* subunit of thiosulfohydrolase; *nar*G: respiratory nitrate reductase 1 ⍰ chain; *amo*A: ⍰ subunit of ammonia monooxygenase) show a clear impact of lugworms. This impact is greatest in the BL where lugworms live and feed (Fig. 3A). Within the PBL, bacterial gene abundances decrease from ∼10^9^ to ∼10^8^ genes g^-1^, archaeal abundances increase from ∼10^6^ to ∼10^7^ genes g^-1^, and eukaryotic abundances decrease from ∼10^8^ to ∼10^6^ genes g^-1^. Similar trends hold in the PL, yet with, on average, ∼3-fold higher bacterial and archaeal and ∼3-fold lower eukaryotic gene abundances. In the BL, prokaryotic and eukaryotic gene abundances decline by ∼2 and ∼1 orders of magnitude, respectively, before increasing again in the UL. rRNA gene abundances decrease less over the interval from 12.5 to 25 cm in controls and show a more continual decrease to the deepest sample (30 cm). Notably, the equally non-bioturbated deepest samples have similar gene abundances in both treatments.

**Fig. 3.**
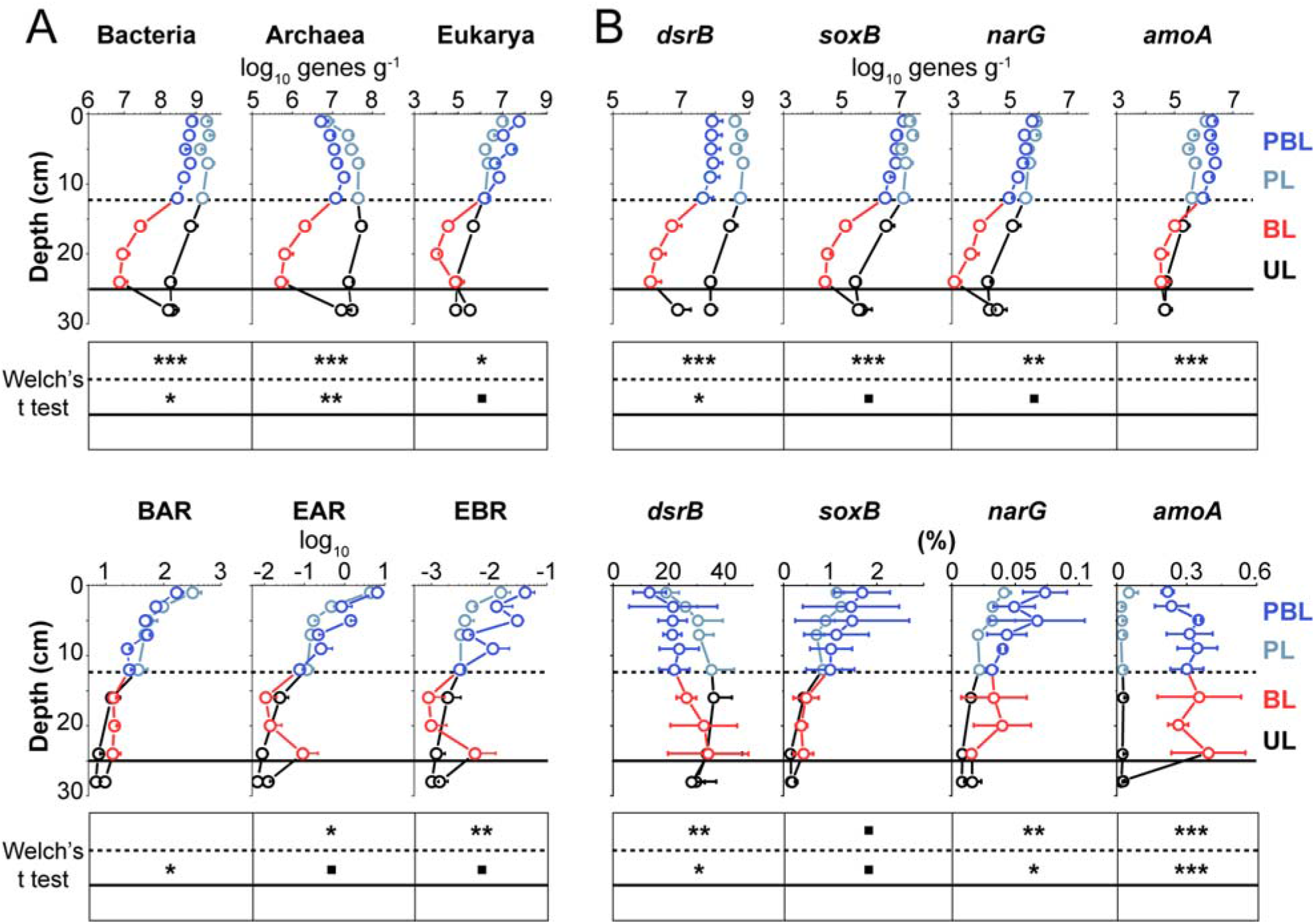
Depth profiles of bacterial and archaeal 16S rRNA gene, eukaryotic 18S rRNA gene, and S- and N-cycling catabolic gene copies in sediments with and without lugworm bioturbation. (A) Depth profiles of bacterial and archaeal 16S and eukaryotic 18S rRNA gene copies (as log10 gene copies g^-1^ wet sediment, upper row), as well as Bacteria-to-Archaea (BAR), Archaea-to-Eukarya (AER), and Bacteria-to-Eukarya (BER) rRNA gene ratios (lower row). (B) Depth profiles of S- and N-cycling catabolic gene copies (sulfate reduction: *dsrB*; sulfide oxidation: *soxB*; nitrate reduction: *narG*; ammonium oxidation: *amoA*; as log10 gene copies g^-1^ wet sediment, upper row), and relative abundances of these genes in %, calculated by dividing *dsrB, soxB, narG*, and *amoA* gene copies by corresponding total 16S rRNA gene copy numbers (lower row). All values represent averages from four plots that were sampled at different time points (error bars denote standard deviations). Significance level of Welch’s t test: \*\*\**p*<0.001, \*\**p*<0.01, \**p*<0.05, · 0.05<*p*<0.1. [Abbreviations: PBL=physically and biologically impacted layer, PL=physically impacted layer, BL=biologically impacted layer, UL=undisturbed layer.]

Gene abundance ratios between domains, calculated as a proxy for relative abundance changes (Fig. 3A), vary with depth in all treatments. Bacteria:Archaea Ratios (BARs) decrease from ∼10^2^ to ∼10^1^, with slightly higher values in the BL than at comparable depths in controls. Eukarya:Archaea Ratios (EAR) and Eukarya:Bacteria Ratios (EBR) decrease from ∼10^1^ to ∼10^−2^ and 10^−2^ to ∼10^−3^, respectively, and are on average 5- and 8-fold higher in the PBL than in the PL, respectively. Notably, EARs and EBRs in worm treatments have localized subsurface peaks (∼16-25 cm) due to increases in eukaryotic gene copy numbers in the lower part of the BL.

Genes involved in reductive (*dsrB*) and oxidative (*soxB*) S cycling are generally 1-3 orders of magnitude more abundant than genes involved in reductive (*narG*) or oxidative (*amoA*) N cycling (Fig. 3B). Depth trends are similar to those in 16S rRNA gene copy numbers. In bioturbated sediments gene abundances are highest in the PBL, followed by a strong decline in the BL, and a slight increase in the UL. In controls, gene abundances decrease less with depth. In both surface sediments (0-12.5 cm) and deeper layers (12.5-25 cm), *dsrB, soxB*, and *narG* copy numbers are lower in worm treatments than in controls. By comparison, *amoA* gene copies are higher in surface sediments of worm treatments, and do not differ significantly between treatments from 12.5-25 cm. Ratios of S- and N-cycling to total 16S rRNA gene copies suggest higher variability in relative abundances of S- and N-cycling microorganisms in bioturbated sediments (Fig. 3B). Overall, ratios are lower (*dsrB*), slightly higher (*soxB*), or clearly higher (*narG* and *amoA*) in bioturbated sediments compared to controls.

### Impact of lugworm bioturbation on benthic community structure

The impact of bioturbation on relative abundances of microbial and eukaryotic taxa is strongest in the BL, where worms mainly live and feed (Fig. 4A). In the following text we present major groups at the class and phylum level with mention of their dominant subgroups in parentheses.

**Fig. 4.**
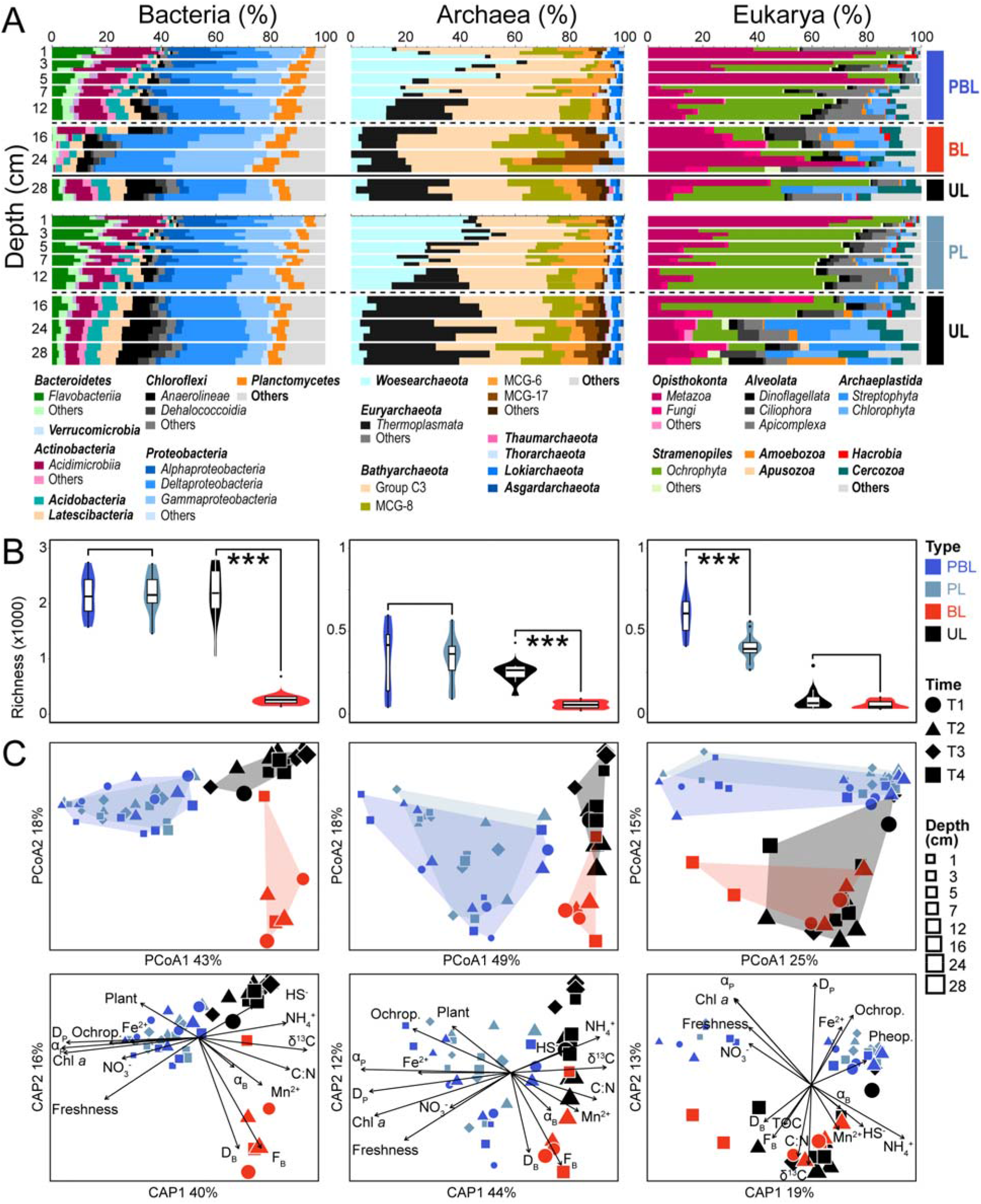
Community analyses on Bacteria, Archaea, and Eukarya in intertidal sediments with and without lugworm bioturbation. (A) Relative abundances of dominant taxa at the phylum or class level. (B) Boxplots of richness across different treatments and sediment layers. Asterisks denote significant differences (*p*<0.001) based on Welch’s t test. Community structuring based on (C) Weighted Unifrac distance-based Principal Coordinates Analysis and (D) Canonical Analysis of Principal Coordinates in relation to environmental variables. All calculations in (B)-(D) were done based on zero-radius operational taxonomic units. Only variables that were significantly correlated with community compositions (PERMANOVA *p*<0.05) are shown in (D). [Abbreviations: PBL=physically and biologically impacted layer, PL=physically impacted layer, BL=biologically impacted layer, UL=undisturbed layer.]

Microbial communities in surface sediments of both treatments are dominated by *Flavobacteriia* (*Flavobacteriaceae*), *Acidimicrobiia* (OM1_clade), *Alphaproteobacteria* (*Rhodobacteriaceae*), *Gammaproteobacteria* (*Woeseiales, Halieaceae*), and *Woesearchaeota* (unclassified). These groups, which we refer to as “surface lineages” from here on, gradually decrease with depth, whereas “subsurface lineages” *Latescibacteria* (unclassified), *Chloroflexi* (*Anaerolineae, Dehalococcoidia*), and archaeal *Thermoplasmata* (Marine Benthic Group D) show the opposite trend. Other groups, such as bacterial *Deltaproteobacteria* (*Desulfobulbaceae, Desulfobacteracea*) and *Planctomycetes* (*Planctomycetaceae*), or archaeal *Bathy-, Thor-, and Lokiarchaeota* show no systematic changes with depth. Sediments of the BL are an exception in that most “surface lineages” and “subsurface lineages” occur at lower percentages than at corresponding depths in controls, while percentages of *Gammaproteobacteria* (*Thiotrichaceae, Piscirickettsiaceae, Ectothiorhodospiraceae*) and several bathyarchaeotal subgroups (Group C3, MCG-8, MCG-17) are elevated.

Eukaryotic communities in surface sediments are dominated by *Metazoa* and *Ochrophyta*, with higher percentages of *Metazoa* (*Arthropoda, Annelida*) and lower percentages of *Ochrophyta* (benthic diatoms *Gedaniella* sp.) in bioturbated sediments compared to controls. In worm treatments, *Dinoflagellata* (*Dinophyceae*), and *Apicomplexa* (*Eugregarinorida*, obligatory invertebrate parasites) increase in deeper layers of the PBL (7-12.5 cm), while *Streptophyta* (seagrass *Zostera marina*, which forms a meadow at the subtidal end of False Bay) and *Chlorophyta* (macroalgal *Ulvophyceae*, unicellular *Chlorodendrophyceae*) become abundant in the BL. Similar trends occur in controls, except that *Ochrophyta* and to a lesser extent *Apicomplexa* dominate over a larger interval (1-16 cm), whereas *Strepto-* and *Chlorophyta* only become dominant below 16 cm. Notably, relative abundances of *Metazoa* (meiofaunal worms *Nematoda* and *Gnathostomulida*) are significantly elevated in the BL while those of *Ochrophyta* are significantly lower compared to the same depths in controls (both *p*<0.01).

### Assembling mechanisms of benthic community

Richness analyses confirm the strong impact of lugworms on intertidal sediment communities (Fig. 4B). While bacterial and archaeal richness are comparable in the PBL and PL, they are clearly lower in the BL than in the same depths of controls. In contrast, eukaryotic richness is higher in the PBL than in the PL but similar in subsurface layers across treatments.

Principal Coordinates Analysis (PCoA, weighted Unifrac distance that considers both phylogenetic distances between taxa and their read percentages) based on zero-radius operational taxonomic units (ZOTUs) confirms the major role of lugworms in structuring benthic communities (Fig. 4C). While Bacteria and Archaea share similar communities between the PBL and PL (both PERMANOVA *p*>0.05), their communities in the BL differ significantly from those at the same depths in controls (*p*<0.01). The same result is obtained with an unweighted Unifrac algorithm that only considers the phylogenetic distances between communities (Fig. S1). By contrast, eukaryotic communities only show significant treatment-related differences when an unweighted Unifrac distance algorithm is used, indicating that bioturbation more strongly affects low-abundance eukaryotic taxa (Fig. S1).

Canonical Analysis of Principal Coordinates (CAP) was used to identify potential drivers of benthic communities (Fig. 4C). The included variables explain 76% of bacterial, 71% of archaeal, and 60% of eukaryotic community variations. The first axis (CAP1) generally differentiates bacterial and archaeal communities of surface (0-12.5 cm) and subsurface (12.5-30 cm) sediments. Surface communities are correlated with physically induced porewater exchange (α_P_) and particle mixing (D_P_), indicators of reactive OC sources (chl *a*, freshness index, *Ochrophyta* and vascular plant *rbc*L copy numbers), and concentrations of NO_3_^-^ and Fe^2+^. Subsurface communities correlate with concentrations of metabolic end products (NH_4_^+^, HS^-^, Mn^2+^), indicators of terrestrial OC sources (C:N, δ^13^C-TOC), and feeding and reworking activities (F_B_, D_B_). The second axis (CAP2) further differentiates prokaryotic communities in the BL from those in the UL, with feeding and reworking activities (F_B_, D_B_), Mn^2+^ concentrations, and C:N showing stronger correlations with communities in the BL. By contrast, eukaryotic communities fall into three clusters. Communities in the top 5 cm of bioturbated sediments and top 1 cm of controls correlate with α_P_, NO_3_^-^, chl *a*, and freshness. Communities from 7-12.5 cm in bioturbated sediments and 3-12.5 cm in controls correlate with D_P_, Fe^2+^, *Ochrophyta rbc*L, and pheopigment contents. Communities from >12.5 cm correlate with F_B_, D_B_, C:N, TOC, δ^13^C-TOC, Mn^2+^, HS^-^, and NH_4_^+^.

### Modification of organismal networks by lugworm activity

The presence of *A. pacifica* profoundly changes benthic community co-occurrence networks, especially among Bacteria and Eukarya (Fig. 5A), by increasing significant correlations among “surface lineages” and reducing correlations among “subsurface lineages”. Major bacterial classes in controls form two correlation clusters, consisting of “surface lineages” (**C1**) and “subsurface lineages” (**C2**) identified earlier. While the same “surface lineages” also cluster in bioturbated sediments (**B1**), lugworm treatments exhibit an additional cluster (**B2;** mainly *Planctomycetacia, Acidobacteria, Bacteroidetes* VC2.1 Bac22; Fig. 5B). Lugworm activity, moreover, breaks the subsurface cluster **C2** into two separate clusters, **B3** (mainly *Latescibacteria*, Bacteroidetes BD2-2, unclassified *Chloroflexi*) and **B4** (*Anaerolineae, Dehalococcoidia, Aminicenantes*). Archaeal clusters in controls also consist of “surface lineages” (**C3**; mainly *Woesearchaeota, Lokiarchaeota* (beta_subgroup), *Altiarchaeales*) and “subsurface lineages” (**C4**; mainly *Thermoplasmata* and *Bathyarchaeota* subgroups). However, in contrast to Bacteria, both clusters are also present in bioturbated sediments (**A1** and **A2**), varying mainly in that the bathyarchaeotal subgroups MCG-10, -14, -17, -24, and -28 are strongly correlated with each other in controls but not in bioturbated sediments.

**Fig. 5.**
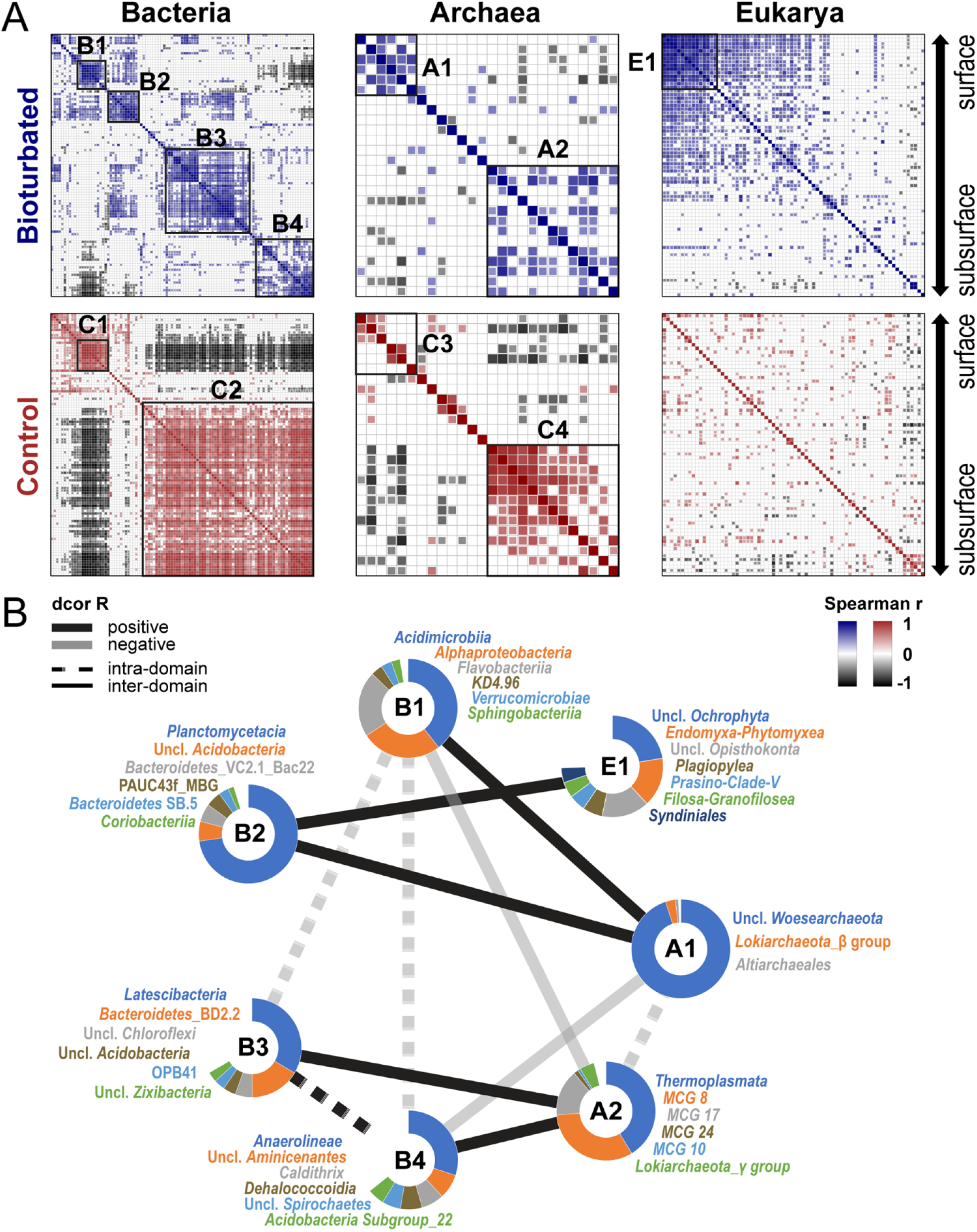
Correlation network analyses of bacterial, archaeal, and eukaryotic communities in bioturbated and control sediments. (A) Pairwise Spearman correlations at the class-level within each domain. Classes are shown in the same vertical and horizontal order for bioturbation treatments and controls. Clusters (B1-4, A1-2, E1, C1-4) were determined based on hierarchical cluster analyses. (B) Intra- and inter-domain relationships of dominant correlation clusters from (A) in bioturbated sediment, calculated based on distance correlations with 999 bootstraps. Only significant correlations (*p*<0.05) are shown in (A) and (B).

Differences in taxa correlations are most pronounced in Eukarya, where *A. pacifica* treatments have many more positive correlations between taxa than controls (Fig. 5A). Strongly correlated eukaryotic groups include certain microalgae (diatoms, *Aurearenophyceae*), protists (*Breviatea, Imbricatea*), and ribbon worms (*Nemertea*). Within these groups, a subset of taxa (**E1**), consisting mainly of *Endomyxa-Phytomyxea, Opisthokonta*, and *Plagiopylea*, show exceedingly strong pairwise correlations (average Spearman *rho*>0.8).

Additional, inter-domain correlation analyses on bioturbated sediments show that surface sedimentary clusters of Bacteria (**B1, B2)** and Archaea **(A1)** are positively correlated, while the eukaryotic cluster **E1** only correlates with bacterial cluster **B2** (all distance correlation R≥ 0.6, *p*<0.01; Fig. 5B). The prokaryotic subsurface clusters **B3, B4**, and **A2** are also positively correlated with each other, and generally negatively correlated with surface clusters. By comparison, controls only show correlations between bacterial and archaeal surface (**C1** and **C3**) and subsurface clusters (**C2** and **C4**, Fig. S2).

## Discussion

Despite being subjected to strong physical forcing by wave action, tides, and daily fluctuations in temperatures, intertidal sediments are rich in sediment macrofauna and represent global hotspots of terrestrial and marine OC cycling (12, 17-22). While it remains unclear how physical forcing and macrofaunal populations compare in their impact on intertidal ecosystem functioning, a diminished importance of bioturbation has been proposed for sediments where hydrological processes dominate (5, 26). Our study shows that, even in intertidal sands that experience strong physical forcing, benthic macrofauna – in this case the lugworm *A. pacifica* – have a major impact on sediment ecosystem functioning.

While modeled rates of physical and biological processes suggest dominance of physical over biological processes in the top few centimeters of sediments (Figs. 1 and 2), a strong impact of bioturbation is evident in deeper sediments that are sheltered from physical forcing (Figs. 3 and 4). Here lugworms significantly alter the porewater chemistry through ventilation. This introduces seawater-derived electron acceptors, e.g. O_2_ and NO_3_^-^, while removing end products of anaerobic mineralization, e.g. NH_4_^+^ and HS^-^. In addition, selective feeding on fine, algal organic matter-rich particles in deeper layers induces a downward movement of sediment, including fresh algal organic matter, and strongly enriches coarse sediment grains and organic macrostructures below burrows. This selective feeding greatly lowers the abundance and richness of, presumably fine particle-associated, Bacteria and Archaea, and negatively affects all but a subset of microbial taxa involved in oxidative N- and S-cycling, macrofaunal symbioses, and degradation of organic macrostructures. Lugworm feeding also negatively impacts eukaryotic abundances and transforms organismal networks, mainly by enhancing network connectivity among “surface lineages” and reducing correlations among “subsurface lineages” (Fig. 5). This strong impact on organismal networks suggests that lugworms fundamentally alter the functioning of intertidal ecosystems. We synthesize the most important findings of our study as described above in a conceptual plot (Fig. 6) and provide detailed explanations in the following sections.

**Fig. 6.**
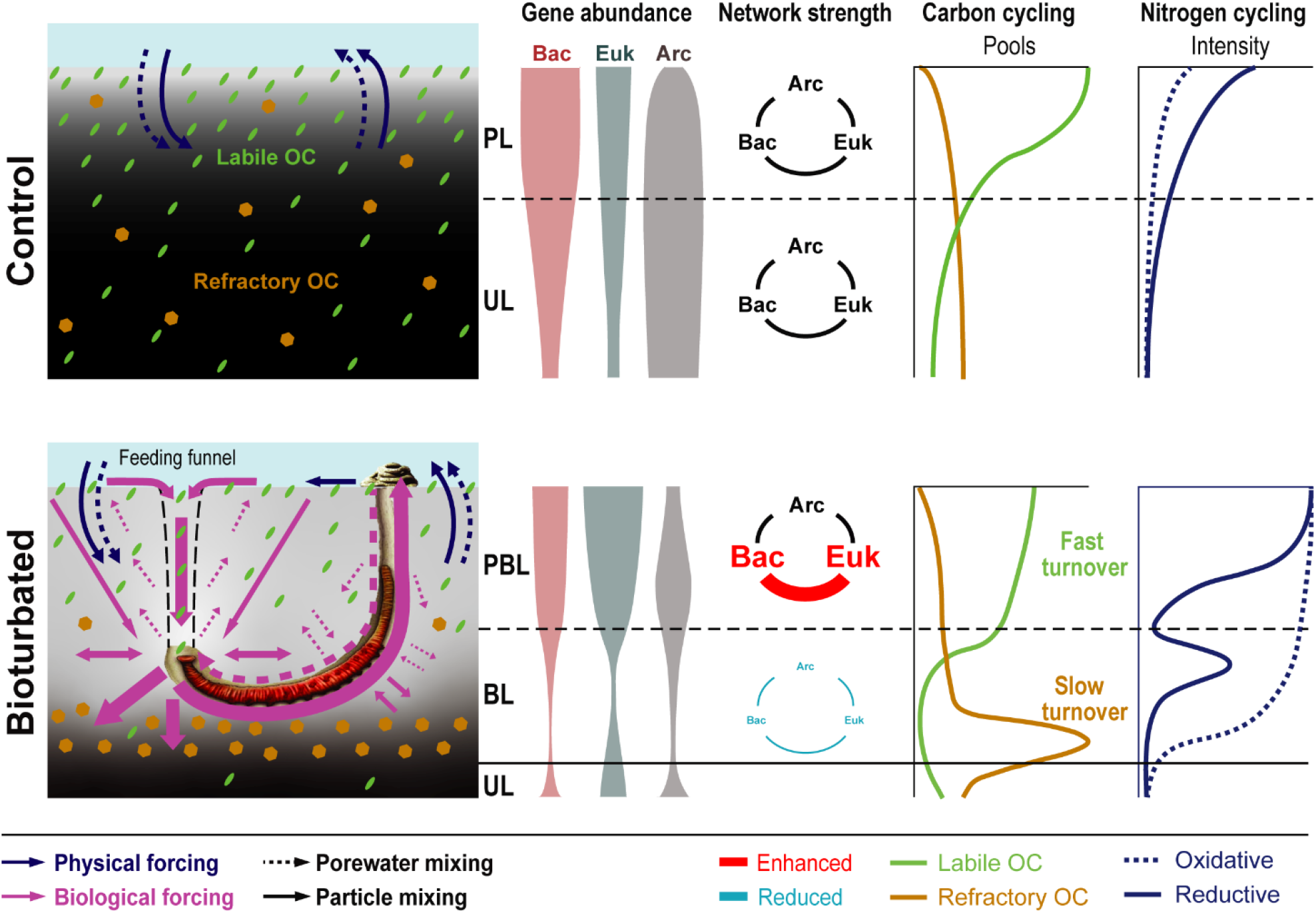
Conceptual diagram comparing ecosystem functioning in lugworm-free controls to lugworm-inhabited intertidal sediment (based on Figs. 1-5). Leftmost panels show differences in porewater and particle mixing between controls and bioturbated plots. In controls, physical forcing dominates, causing porewater and particle mixing to be mainly limited to surface sediments (PL). In lugworm treatments, worm activity in subsurface sediments profoundly impacts porewater and particle mixing throughout. This results in strong differences in redox gradients, whereby lugworms increase the depth of oxidizing conditions (light gray background color) and consequent onset of reducing conditions (black color). The biggest impact of worms is in the BL, where worms spend most of their time in the horizontal section of their burrows. High rates of sediment ingestion, whereby worms selectively feed on labile OC-rich fine particles, effectively drives down bacterial, archaeal, and eukaryotic populations. Simultaneously, worm feeding avoidance of (mainly terrestrial) macrostructures increases the burial of refractory OC. In sediments overlying burrows (PBL), lugworm activity strongly promotes intra- and inter-domain bacterial and eukaryotic network connectivity, while the opposite is the case in the BL where worms feed. By contrast, lugworm activity only has a minor impact on intra-domain archaeal networks, or archaeal networks with other domains. Finally, bioirrigation by lugworms alters the distribution of oxidative and reductive N- (and to a lesser extent S-) cycling, enhancing the relative importance of ammonium oxidation (nitrification) and nitrate reduction (denitrification) throughout bioturbated sediment. [Abbreviations: PBL=physically and biologically impacted layer, PL=physically impacted layer, BL=biologically impacted layer, UL=undisturbed layer.]

### Physical processes mainly control surface sedimentary communities

Bacterial and archaeal communities in surface sediments (<10 cm) are distinct from those in other sediment layers (Fig. 4), but do not change significantly in the presence of *A. pacifica* (PERMANOVA *p*>0.05). Though physical and biological processes both enhance porewater exchange and sediment mixing, our data indicate that physical ventilation (α_P_) and particle mixing (D_P_) dominate in surface sediments (Figs. 1 and 2). These results are in line with those of a tracer-based study conducted in a nearby but lower part of the False Bay intertidal area, that reported physical porewater advection throughout the upper 10 cm of sediment (27). α_P_ and D_P_ are the variables that explain the highest percentages of bacterial (∼30%, ∼28%) and archaeal (∼33%, ∼30%) community variation in surface sediments (Fig. 4C), whereas α_B_ and D_B_ appear more important in the BL, where both reach their peak values.

Physical factors appear to control microbial communities in surface sediments through their strong impacts on sediment geochemistry. α_P_ and D_P_ are significantly positively correlated with NO_3_^-^ concentrations, chl *a* content, OC freshness, and *rbc*L copy numbers (especially *Ochrophyta*), and significantly negatively correlated with C:N, δ^13^C-TOC, HS^-^ and NH_4_^+^ (Fig. S3A). All of these variables correlate significantly with the bacterial and archaeal community structure in surface sediments (CAP, Fig. 4C), indicating that physical processes drive microbial communities in surface sediments through increased input of seawater-derived electron acceptors, removal of potentially inhibitory or toxic (anaerobic) mineralization end products, and transport of labile, e.g. benthic *Ochrophyta* (diatom)-derived, OC from the sediment surface to deeper layers. While the geochemical agents are similar to those previously proposed to drive microbial community structure in continental shelf surface sediments (16), it is physical forcing rather than bioturbation that controls these variables in intertidal surface sands of False Bay.

Matching the fluctuating physical and redox conditions in sandy, advection-controlled surface sediments (28, 29), phylogenetic analyses indicate high metabolic flexibility and/or physiological resilience among “surface lineages”. Dominant “surface lineages”, such as *Flavobacteriaceae* (*Flavobacteriia*) and *Rhodobacteraceae* (*Alphaproteobacteria*), are widespread across diverse habitat types (seawater, sediment, soil, biofilms) and include (facultatively) aerobic and anaerobic chemoorgano- and photo(organo)trophic members (30, 31). Other dominant “surface lineages” can switch between aerobic respiration and fermentation during redox oscillations (*Woeseiales* (*Gammaprotebacteria*) (29)), scavenge oxygen for anaerobic growth (*Desulfobulbaceae* (*Deltaproteobacteria*) (32)), and occur widely in oxic and anoxic surface sediments (*Woesearchaeota* (33)).

Despite the likely dominance of physical over biological processes, bioturbated surface sediments have a higher eukaryotic abundance and richness, and different eukaryotic community structure than controls (Figs. 3, 4). Of all sequenced eukaryotic ZOTUs, 44% are shared between the PBL and PL, while the rest are unique to the PBL (35%) or PL (21%). Dominant unique taxa in bioturbated sediments include many meiofauna (Fig. S4), such as parasitic worms (*Chromadorea* within *Nematoda* (34)) and alveolates (*Actinocephalidae* within *Gregarinomorphea* (35)), and worms that feed on microbial populations in suboxic, sulfur-rich environments, often near polychaete burrows (*Haplognathia* within *Gnathostomulida* (36)). In addition, relative abundances of surface deposit-feeding polychaetes (*Boccardiella* spp. (37)) were clearly elevated in the PBL (Fig. S5) and explain the high percentage of *Metazoa* at 5 cm sediment depth in bioturbated sediment (Fig. 4A).

Enhanced sediment permeability leading to increased input of oxygenated seawater and removal of toxic metabolites, e.g., HS^-^, in bioturbated surface sediments may promote the thriving of these meiofauna and other eukaryotic organisms (22, 36), as could higher energy supply due to enhanced input of labile OC. The latter explanation is supported by higher freshness values and lower pheopigment contents in the PBL compared to the PL (Fig. 2B). Higher input and turnover of labile OC likely results from subsurface “conveyor feeding” by *A. pacifica*, whereby the microalgae-rich surface sedimentary layer is perpetually “subducted” into deeper layers via worm feeding funnels (Fig. 6). The subsequent return of ingested sediment to the surface as nutrient-rich feces, and the upward advective flow of nutrient-rich porewater from depth, could foster rapid regrowth of microalgae and lead to enhanced biomass turnover in worm-inhabited sediments (38).

Yet, the reason why higher input and turnover of labile OC, and thus bioavailable energy, in the PBL would increase eukaryotic biomass but not that of Bacteria and Archaea (Fig. 3A), is unclear. Past studies have shown that worm feces are microbially recolonized from surrounding sediments within hours (14) and support high growth rates that lead to recovery of microbial communities within 24 hrs (39). We speculate that motile, grazing meiofauna, such as *Gnathostomulida*, which actively move through the interstitial space of sandy sediments (36), maintain their vertical position even when exposed to a downward movement of surrounding sediment particles. This may enable these meiofauna to exploit the enhanced supply of microbial biomass in the PBL while at the same time avoiding predation by *A. pacifica* in the underlying BL. The same would not apply to most Bacteria or Archaea, which are particle-attached and grazed upon by motile meiofauna during transport through the PBL and subsequently by *A. pacifica* in the BL. Exceptions might include free-living prokaryotes, e.g. ammonium-oxidizing archaea (40), which occur at an order of magnitude higher abundances in the PBL than the PL (Fig. S6). These archaea could evade ingestion by particle-feeding *A. pacifica*, and benefit from the elevated supply of O_2_ from bioirrigation in sediments above *A. pacifica* burrows, as indicated by the significant positive correlation between *amo*A copies and α_B_ (Fig. S3B).

### Lugworm activity drives subsurface communities

Worm feeding (F_B_) and sediment mixing (D_B_) have a strong impact on the community structure of Bacteria, Archaea, and Eukarya in subsurface sediments (Fig. 4C), and both negatively affect prokaryotic gene copy numbers, and prokaryotic and eukaryotic richness (Figs. 3, 4B, and S3B).

While D_B_ values are positively correlated with freshness index, in line with increased input and cycling of labile OC in bioturbated sediments, F_B_ values have positive correlations with C:N (Fig. 2; Fig. S3A). Pheopigment contents and *rbc*L copy numbers are furthermore more strongly (negatively) impacted by F_B_ than by D_B_ (Fig. 2; Fig. S3A). These correlations are in line with the observation that *A. pacifica* selectively feeds on microalgae- and prokaryote-rich fine particles (13). This leads to the gradual enrichment of non-ingestible coarse sands (Fig. 1A) and macrostructures, such as woody debris, seaweed detritus, and bivalve shells below lugworm burrows (‘biogenic bedding’), as observed for other ecosystems inhabited by deposit-feeding fauna (4, 41). This selective feeding has a striking impact on bulk OC compositions, as evidenced by C:N and δ^13^C-TOC. These range from fresh microalgae-dominated surface sediments (C:N: 7-10; δ^13^C-TOC: -23 to -18‰) to OC compositions that are indicative of mainly terrestrial C3 vascular plant origin (C:N: ∼30; δ^13^C-TOC: ∼-27‰; (42)) below lugworm burrows (Fig. 2). The low *rbc*L copy numbers of vascular plants (Fig. 2B) are consistent with this vascular plant matter being dominated by wood and/or highly degraded detritus. While downward mixing of fresh OC, bioirrigation, and release of nutrient-rich secretes were previously shown to stimulate growth of meio- and microorganisms within burrows of *A. pacifica* and other sediment macrofauna (“microbial gardening” (13, 43)), our study suggests that these effects are offset in bulk sediments by feeding-induced negative impacts.

The strong impact of worm feeding is also reflected in distinct organismal communities between bioturbated and non-bioturbated subsurface sediments (Fig. 4). Relative abundances of microalgae (mainly benthic diatom *Gedaniella* spp.), algal polysaccharide degraders (Flavobacteriaceae (30)), and the algae-associated OM1_clade (Acidimicrobiia (44)) decrease in subsurface layers, in line with A. pacifica feeding on fine, microalgae-rich particles. Yet, matching the elevated content of wood, seaweed, and shells that are rich in lignin, cellulose, or chitin (45), other lineages increase, including bathyarchaeotal subgroups that have been linked to the anaerobic fermentative or acetogenic degradation of lignin (MCG-8 (46)), cellulose (MCG-8, -15, -17 (47)), and chitin (MCG-15 (Group C3) (47)). These lineages may be protected from lugworm ingestion by being associated with detrital macrostructures.

In addition, subsurface bioirrigation and commensal interactions could play a role in structuring subsurface communities. Percentages of several gammaproteobacterial families (*Thiotrichaceae, Piscirickettsiaceae*, unclassified Ca. *Thiobios*) increase in the BL (*p*<0.05). Members of *Thiotrichaceae, Piscirickettsiaceae*, and Ca. *Thiobios* include sulfide oxidizers that use O_2_ (48, 49) or NO_3_^-^ as electron acceptors (50). These groups may benefit from input of O_2_ and NO_3_^-^ by subsurface bioirrigation, which would match the higher relative percentages of *narG* and *soxB* genes in the BL (Fig. 3B). Intriguingly, members of these families have also been found in associations with marine fauna (*Thiotrichaceae*: echinoids (51); *Piscirickettsiaceae*: fish (48); Ca. *Thiobios*: ciliates (49)). In contrast, relative abundances of “subsurface lineages” such as bacterial Chloroflexi (mainly Anaerolineaceae (52) and Dehaloccocoidia (53)) and archaeal *Euryarchaeota* (mainly Marine Benthic Group D within *Thermoplasmata* (54)), that include anaerobic fermenters and acetogens, decrease in the BL, matching the reported negative impact of bioirrigation on these anaerobic lineages (16).

The metazoan community is dominated by free-living Nematoda (*Trefusidae*) and *Gnathostomulida* (again *Haplognathia*), which both increase in diversity and relative abundances in the BL (Figs. S4, S5). Members of both groups are frequently enriched in suboxic sediments but differ in diets (*Trefusidae*: selective deposit feeders (55); *Haplognathia*: microbial grazers (36)). Both groups may live by scavenging on macrofaunally released O_2_, food scraps, and metabolites, and evade worm predation by means of their free-living lifestyles and/or by inhabiting coarse particles that are not ingested by *A. pacifica* (36, 55, 56).

### Lugworms modify sediment community networks

Our research shows that lugworms strongly alter community networks by increasing positive correlations among surface taxa (**B2, A1, E1**) while reducing those among subsurface taxa (**B3, B4, A2;** Fig. 5). In addition, negative correlations between surface and subsurface bacterial lineages (**C1** vs. **C2**) are greatly reduced in the presence of lugworms.

Worm bioturbation supports unique bacterial and eukaryotic clusters (**B2** and **E1**) among “surface lineages”, which are, moreover, significantly correlated with each other (Fig. 5). The bacterial cluster **B2** contains putative aerobic degraders of algal polysaccharides (*Planctomycetacia*, mainly *Pirellula* and *Rhodopirellula* (57)) and cellulose (Bacteroidetes VC2.1_Bac22 (58)), and metazoan symbionts (PAUC43f_marine_benthic_group (59)). The eukaryotic cluster **E1** includes microalgae (*Ochrophyta*, Prasino-Clade-V), plant pathogens (*Endomyxa-Phytomyxea* (60)), bacterivorous protists (*Filosa Granofilosea* (61)), and ciliates that are known to host denitrifying endosymbionts (*Plagiopylea* (62)). While the metabolisms of these groups are typical of intertidal surface sediments, they are diverse and do not give clear indications that members of **B2** and **E1** enter biological consortia or co-dependencies in the presence of bioturbation. Instead these correlations could also be driven by factors that co-vary in lugworm-inhabited sediments, e.g. higher vertical sediment transport, increased input of labile OC from above and of O_2_ from below, higher meiofaunal and metazoan population size. These same variables, especially the higher rates of vertical sediment transport and bioirrigation, may also reduce the vertical zonation of surface and subsurface bacterial lineages and explain the absence of negative correlations between surface and subsurface bacterial lineages in the presence of lugworms.

Interestingly, presence of lugworms reduces correlations in subsurface sediments compared to *A. pacifica*-free controls. This is most apparent in the bacterial cluster **C2**, which is broken down into two separate smaller clusters (**B3** and **B4**; Fig. 5A). Major lineages of **B3** include putative sulfate reducers (*Zixibacteria* (63)), widespread but poorly known members of the phylum Bacteroidetes (BD2-2), most members of which are primary fermenters of carbohydrates and proteins (64), and members of *Latescibacteria*, which genomic analyses suggest are primary fermenters of algal polysaccharides and glycoproteins (65). **B4** also comprises groups that are linked to the anaerobic degradation of carbohydrates and proteins (*Aminicenantes* (66); *Anaerolineae*, mainly *Anaerolineaceae* (52); *Dehalococcoidia* (53); *Deferribacteres*, mainly *Caldithrix* (67)). Yet, while **B3** member are mostly found in marine sediments and seawater, **B4** members are also widespread in activated sludge, terrestrial aquifers, and terrestrial deep subsurface environments (see references above), where the content of algal detritus is low. It is thus possible that differences in primary OC sources drive the observed patterns. The near absence of **B3** from the terrestrial OC-dominated deep part of the BL matches the fact that most of its members are linked to algal OM degradation (Fig. S7). By the same token, cluster **B4** occurs at high relative abundance in the terrestrial OC-dominated deep layer, and could include key degraders of vascular plant detritus. Thus, the separation of **B3** and **B4** in bioturbated sediment may be the outcome of feeding-related partitioning of OC pools by *A. pacifica*.

## Conclusions

Even though hydrodynamic forcing and macrofaunal bioturbation are the main sources of disturbance in coastal sediment (12, 15-18, 20-22), the relative impact of both processes on biological communities was previously not known. We show that intertidal sand ecosystems dominated by lugworms can be divided into distinct hydrodynamically and biologically controlled layers. Hydrodynamic forcing, by regulating OC and electron acceptor inputs, is the main driver of biological communities in surface sediments. In contrast, communities in deeper layers, that are sheltered from hydrodynamic forcing, are largely controlled by lugworm activities. Though lugworms clearly modify OC and electron acceptor inputs in these deeper sediments, their most important influence on organismal communities is through selective feeding on fine particles. This selective feeding effectively depletes labile microalgal organic matter, drastically lowers abundances of Bacteria, Archaea, and Eukarya, strongly weakens biological networks, and promotes the burial of refractory OC. The clear division into hydrodynamically and bioturbation-controlled vertical zones provides a basis for integrating physical and biological factors into ecological and biogeochemical models of globally distributed intertidal sand ecosystems.

## Materials and methods

### Study area

The study area is located in the northwest part of the False Bay Biological Preserve on San Juan Island, USA (Fig. S8). False Bay is an intertidal embayment (∼1 km^2^), that is covered by coarse sand at the mouth and finer silty and muddy sand along the margins, and is subject to mixed semi-diurnal tides (68). *A. pacifica* dominates several upper intertidal areas of False Bay, in some places in close proximity with two species of ghost shrimp (Thalassinidea; also see 27). We here focus on an area where ghost shrimp or other large macrofauna besides *A. pacifica* were absent at the time of the study. This area is located 150-200 m east and ∼500 m south of where two small streams enter the bay. Both streams were nearly dried up during the sampling period.

### Site descriptions

To determine the impact of hydrodynamic processes and bioturbation on sediment geochemistry and biological communities in intertidal sediments, we compared bioturbated sediment dominated by *A. pacifica* (30-40 individuals m^-2^) to an adjacent control area (Fig. S8). This control area was nearly macrofauna-free, but had otherwise similar hydrological conditions, lithologies, and OC sources. Throughout the study period, sediment porewater in both areas had chloride (Cl^-^) concentrations (470-510 mM) and salinities (∼30-33 psu; Fig. S9) in the range of seawater from the Strait of Juan de Fuca (30-34 psu, 69), indicating minimal groundwater seepage. Temperatures during the day varied from 17 to 27 °C at the sediment surface and decreased to steady values of 15-17 °C below ∼15 cm (Fig. S10). Sites were exposed for 6-8 hours during each low tide, for the duration of which sediments remained water-saturated to the sediment surface.

### Study Design

To maximize comparability between sampling dates, all sampling was initiated within 1 hour before to 2 hrs after tidal exposure [T1: August 5^th^, ∼2 hrs after exposure; T2 (August 19^th^) and T3 (August 27^th^): 1 hour before exposure with 10-20 cm of water; T4 (September 7^th^): ∼1 hr after exposure]. All sampling and microsensor measurements were completed within ∼1 hr.

Sampling of natural sediments was complemented by field manipulation experiments in which surface sediments were defaunated by sieving through a 1-mm mesh, homogenized, returned to the original area, pre-incubated for 3 days, and subsequently refaunated with *A. pacifica* or kept defaunated. Sampling was performed at the same dates as for field investigations. These experimental plots were used to quantify bioirrigation activity (as explained later).

### Microsensor profiling

Depth profiles of dissolved O_2_, HS^-^, pH, and redox potential were determined at all sampling plots using a field micro profiling system equipped with Clark-type microsensors (tip size: 200 μm, Unisense, Denmark).

### Sampling

Sediment cores (∼40 cm) were taken using 90-mm diameter plexiglass core liners on bulk sediments. Porewater was sampled using rhizons (0.2 μm pore size, Rhizosphere, Netherlands), that were inserted through pre-drilled holes in the liner, and treated with acid or base for cation or anion analyses, respectively (for details see (70)). All porewater samples were immediately stored inside coolers with -20°C ice packs. After porewater sampling, sediments were extruded and sliced into 1-2 cm thick vertical sections. Samples for microbiological and solid-phase geochemical analyses were taken from the center of each section using sterile, cut-off 5 mL syringes, and immediately frozen in coolers with dry ice.

### Solute-phase geochemical measurements

Dissolved Fe^2+^ and Mn^2+^ concentrations were measured by Inductively Coupled Plasma-Optical Emission Spectroscopy (ICP-OES 5100, Agilent Technologies (70)). Concentrations of NH_4_^+^ (71), NO_3_^-^ and NO_2_^-^ (72), and HS^-^ (73) were determined photometrically on a plate reader (Synergy HT, BioTek). Concentrations of SO_4_^2-^ and Cl^-^ were measured by Ion chromatography (DIONEX DX-320 (70)). Standards consisted of solutions of MilliQ water containing analytical-grade ammonium chloride, zinc sulphide, and sodium salts of sulfate, chloride, nitrate, and nitrite, and an ICP-multielement standard solution (all Sigma-Aldrich, Switzerland).

### Solid-phase geochemical measurements

Chl *a* and its degradation products (pheo-pigments) were extracted from ∼1.5 g wet sediments using acetone and quantified spectrophotometrically using an acidification protocol ((74), Cary 50, UV-Vis, Varian). Chl *a* freshness indices were calculated based on ratios of Chl *a* to the sum of Chl *a* and pheo-pigments (75). Total organic carbon (TOC), total nitrogen (TN), and their isotopic compositions (δ^13^C-TOC, δ^15^N-TN) were measured on decarbonized samples by elemental analyzer/isotope ratio mass spectrometry (16). Ratios of TOC to TN (C:N) served as indicators of OM source and/or quality (42).

### DNA and RNA extraction

Total DNA and RNA was simultaneously extracted from ∼0.2 g of wet sediment following lysis protocol I of Lever et al. (76). This protocol involves chemical (phenol-chloroform-isoamylalcohol, lysis solution I) and mechanical cell lysis (bead-beating: 0.1-mm Zirconium beads), followed by 2× washing with chloroform:isoamyl alcohol (24:1), precipitation with a mixture of linear polyacrylamide, sodium chloride and isopropanol, and purification with the CleanAll DNA/RNA Clean-Up and Concentration Micro Kit (Norgen Biotek Corp., Canada).

### Quantitative PCR (qPCR)

Bacterial and archaeal 16S rRNA genes, eukaryotic 18S rRNA genes, *Ochrophyta rbc*L genes, and functional genes encoding the ammonia monooxygenase subunit A of ammonia-oxidizing Archaea (AOA) and Bacteria (AOB) (*amoA*), respiratory nitrate reductase (*nar*G) of denitrification, dissimilatory sulfite reductase (*dsr*B) of sulfate reduction, and thiosulfohydrolase (*sox*B) of sulfide oxidation were quantified by SYBR-Green based qPCR assays using a LightCycler 480 II (Roche Life Science, Switzerland). Published primer pairs and a newly designed primer set for *sox*B were applied (Tables S1, S2).

### Sequence and bioinformatic analyses

Amplicon libraries were prepared according to (16) using published 16S and 18S rRNA gene primers (Table S1) and by sequencing via the MiSeq platform (Illumina Inc., USA). Raw sequence reads were processed following a bioinformatic pipeline in Deng et al. (16).

### Physically and macrofaunally induced mixing of porewater and sediment particles

Physical and biological porewater mixing was determined by numerical modeling of porewater profiles of DIC and SO_4_^2-^ in defaunated and refaunated sediment plots (see Supplementary Materials for details). Physical and biological porewater exchanges were described as non-local exchanges with depth-dependent physical mixing frequencies *a*_*P*_*(x)* and bioirrigation coefficients *a*_*B*_*(x)*, respectively. Solid-phase mixing was determined by adjusting diffusive mixing coefficients (D_P_, D_B_) to match measured with simulated chl *a* and pheopigment profiles in natural sediments (see Supplementary Materials for details).

### Feeding intensity

Feeding intensity (F_B_), which we define as the selective feeding of lugworms on fine sediment particles, enriches coarse grain sizes within and below the living depths of lugworms (Fig. 1A; 4, 13). We provide a proxy for F_B_ based on differences in sand fractions between worm (*f*_Φsand_) and control treatments (*f*′_Φsand_):

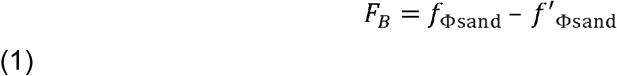

### Multivariate statistical analysis

All statistical analyses were performed in R (77). Microbial richness was calculated based on rarefied datasets of observed ZOTU numbers. Ordination analyses (PcoA, CAP) with weighted Unifrac distances were performed using the ‘phyloseq’ package (78). Permutational multivariate analysis of variance (PERMANOVA, permutations = 999) and statistical tests (two-tailed pairwise t test and Welch’s t test) were performed using the ‘vegan’ and ‘stats’ packages, respectively (79). Community networks were constructed based on the pairwise Spearman correlations between taxa and visualized using the ‘corrplot’ package with the hierarchical clustering method (80). Inter-domain correlation strength was quantified by calculating the coefficients of distance correlations (dcor R) using the ‘energy’ package (81).

## Acknowledgments

This study was funded by Swiss National Science Foundation Project 205321_163371 (M.A.L.). C.M. was funded by National Science Foundation Long-Term Ecological Research program NSF OCE-1832178 (C.M.). We thank Bernadette Holthuis, Jeannie Meredith, Megan Dethier, Michelle Herko, Stacy Markman (Friday Harbor Laboratories, University of Washington) for administrative and scientific support during the sampling campaign at the False Bay Biological Preserve, Iso Christl and Ruben Kretzschmar (ETH Zurich) for technical and instrument support with the analysis of dissolved metals, Madalina Jaggi (ETH Zürich) for assistance with carbon and isotopic analyses, and Jean-Claude Walser, Silvia Kobel, and Aria Minder (Genetic Diversity Centre, ETH Zurich) for llumina sequencing and bioinformatic support.

