## Supplementary materials for "Deposit-feeding worms control subsurface ecosystem functioning in intertidal sediment with strong physical forcing"

Present addresses:

**This PDF file includes:**

Supplementary Text

Figs. S1 to S11

Tables S1 to S2

**Supplementary Text**

**Intertidal manipulation experiments for the quantification of porewater mixing profiles by physical and macrofaunal processes**

Sediment porewater exchanges with overlying seawater by physical mixing (e.g., tidal flushing, wave action) and by bioirrigation were determined based on *in situ* manipulation experiments. The top 30 cm of a 2×3 m^2^ large intertidal area were defaunated by sieving through a 1-mm mesh, thoroughly homogenized, and returned to the original area. After pre-incubation for 3 days, this defaunated area was divided into two equally sized parts by inserting a nylon-mesh barrier through the middle. One half was kept defaunated (**NF**), while the other half was refaunated by adding *Abarenicola pacifica* at natural densities (**AP**, 35 individuals m^-2^). NF treatments were subject to the influence of physical mixing only, while AP treatments were subject to both physical mixing and lugworm bioirrigation. Porewater sampling in the manipulation plots was performed over a period of two weeks, on the same dates as sampling for field investigations on natural sediments, to ensure that modelled rates were subject to similar impacts of temperature, wave action, and tidal movements.

**Modeled rates of porewater exchange by physical forcing and bioirrigation**

Porewater exchanges due to physical mixing ($\alpha_{P}$) and bioirrigation ($\alpha_{B})$were defined as the non-local transport of fluids between sediments and overlying water as a function of sediment depth ($\alpha_{P}$*(x)*, $\alpha_{B}$*(x))*:

$\phi\frac{\partial C}{\partial t}=\frac{\partial}{\partial x}\left( \phi D\frac{\partial C}{\partial x} \right)+R+\alpha_{P}\phi\left( C_{0}-C \right)+\alpha_{B}\phi\left( C_{0}-C \right)$	(S1)

where *φ* is porosity (set to 0.65), *D* is the *in situ* diffusion coefficient corrected for tortuosity, $D={D_{mol}}/\left( 1-2ln\left( \phi\right) \right)$ (1), *R* is the species-specific net reaction rate, and *C_0_* is the concentration in overlying water. Changes in physical porewater mixing intensity with depth were represented by the function $\alpha_{P}(x)={\alpha_{P,0}}/\left( 1+e^{\alpha_{P,1}\left( x-\alpha_{P,2} \right)} \right)$, which reflects the fact that physical mixing is the result of external forcing (e.g. wave) from overlying water. The reaction rate profile was assumed to vary monotonously with depth, $R(x)={r_{0}}/\left( 1+e^{r_{1}\left( x-r_{2} \right)} \right)$. Such a profile is able to – with a minimum of adjustable parameters – reflect high DIC production from the mineralization of newly added reactive organic matter at the sediment surface, while capturing decreases in organic matter mineralization rates with depth (e.g. 2).

*A. pacifica* are deposit feeders that live in J-shaped burrows and regularly pump seawater into sediments inducing porewater movement (3). The effect of lugworm pumping on porewater movement has been shown to be most pronounced near and above the base of the feeding funnel (injection site), with porewater flow spreading (and hence slowing decreasing) away from this location towards the sediment water interface. Porewater advection below the injection site is limited (4). This is captured by a bioirrigation profile that is maximal at the approximate burrow depth, and decreases rapidly below the burrow and more gradually upwards to the sediment-water interface (4).

The parameterization of rate *R(x)*, physical mixing coefficients $\alpha_{P}$*(x)*, and bioirrigation coefficients $\alpha_{B}$*(x)* was established by simulating the DIC concentration profiles. First, physical mixing and net DIC production profiles were estimated in the NF treatment (i.e. $\alpha_{B}$ = 0; *R(x)* and $\alpha_{P}$*(x)*). Using the initial DIC profile as starting conditions, the parameters were adjusted to fit the DIC concentration profile measured 2 weeks later. Bioirrigation intensities representative of a bioturbated environment were then determined from *A. pacifica* (AP) treatments, assuming the same physical mixing coefficient profile across NF and AP plots. We also assumed the same DIC production profile in NF and AP plots, as the sediments were homogenized prior to both treatments and thus had similar and homogeneous starting distributions of reactive organic matter (5). The parameterization we applied resulted in generally good fits between measured and modeled porewater concentration profiles (Fig. S10).

**Modeled rates of sediment mixing by physical and biological forcing**

The timescale (2 weeks) of the manipulation experiments turned out to be too short to observe significant changes in distributions of sediment particles. Solid-phase mixing was thus estimated based on vertical profiles of chlorophyll *a* (chl *a*) and pheopigment contents in natural sediments of bioturbated and control sites (as shown in Fig. 2). In the bioturbated sediments, rates of total sediment mixing (D_T_) include mixings from both physical (D_P_) and biological mixings (D_B_), while in the control sediments, D_T_ includes mainly physical mixing (D_P_). Sediment mixing was thus modeled as:

$\frac{\partial C}{\partial t}=\frac{\partial}{\partial x}\left( D_{T}\frac{\partial C}{\partial x} \right)-k_{C}C$	(S2)

$\frac{\partial P}{\partial t}=\frac{\partial}{\partial x}\left( D_{T}\frac{\partial P}{\partial x} \right)+k_{C}C-k_{P}P$	(S3)

$D_{T}=D_{B}+D_{P}$	(S4)

where C is the content of chl *a*, P is the content of pheopigments, k_C_ and k_P_ are the rate constants for the breakdown of chl *a* and pheopiments, respectively. k_C_ is set to 0.04 d^-1^ (average value from different literature sources 6-8). D_T_ is a mixing coefficient for total mixing, D_P_ and D_B_ are the mixing coefficients for physical and biological mixings, respectively. Given the negligible impact of macrofaunal bioturbation in control sediments, D_B_ was set to zero to solve the term for D_P_ in these (equation S4). The same D_P_ was then applied to bioturbated sediments to solve for D_B_.

We divide the sediment column into three layers (0-12.5, 12.5-25 and >25 cm; see Results of main text for details), each with a constant value of D_T_. The resulting analytical solutions for the concentration profiles in layer *i* are

$C\left( x \right)=M_{1,i}exp\left( \sqrt{{k_{C}}/{D_{T,i}}}x \right)-M_{2,i}exp\left( -\sqrt{{k_{C}}/{D_{T,i}}}x \right)$	(S5)

and

$P\left( x \right)=N_{1,i}exp\left( \sqrt{{k_{P}}/{D_{T,i}}}x \right)-N_{2,i}exp\left( -\sqrt{{k_{P}}/{D_{T,i}}}x \right)+\frac{k_{C}}{k_{P}}C$	(S6)

The constants M_j,j_ and N_j,i_ were determined by the boundary conditions including the measured concentrations at the top (1 cm), depths between layers (i.e. 12.5 and 25 cm), flux continuation, and no-gradient conditions at infinite depth. We applied a k_P_ value of 0.015 d^-1^ that was measured in sandy sediments of an adjacent coastal location (Dabob Bay, Washington, USA; 9) for the control treatments. For the bioturbated treatment, however, this relatively low k_p_ did not reflect the fast removal and consequently low concentrations of pheopigments. A higher k_p_ value of 0.03 d^-1^ that approximates the rate constant of chl *a* was therefore applied for bioturbated treatment (value measured in coastal sandy/silty sediments that were subject to hydrodynamic forcing and bioturbation, Long Island Sound; 7). We then optimized the D_T_ value in each layer *i*. The parameterization we applied generally resulted in good fits for all data of chl *a*, pheopigments, as well as freshness index across the four different time points (Fig. S11).

**Supplementary Figures and Tables**

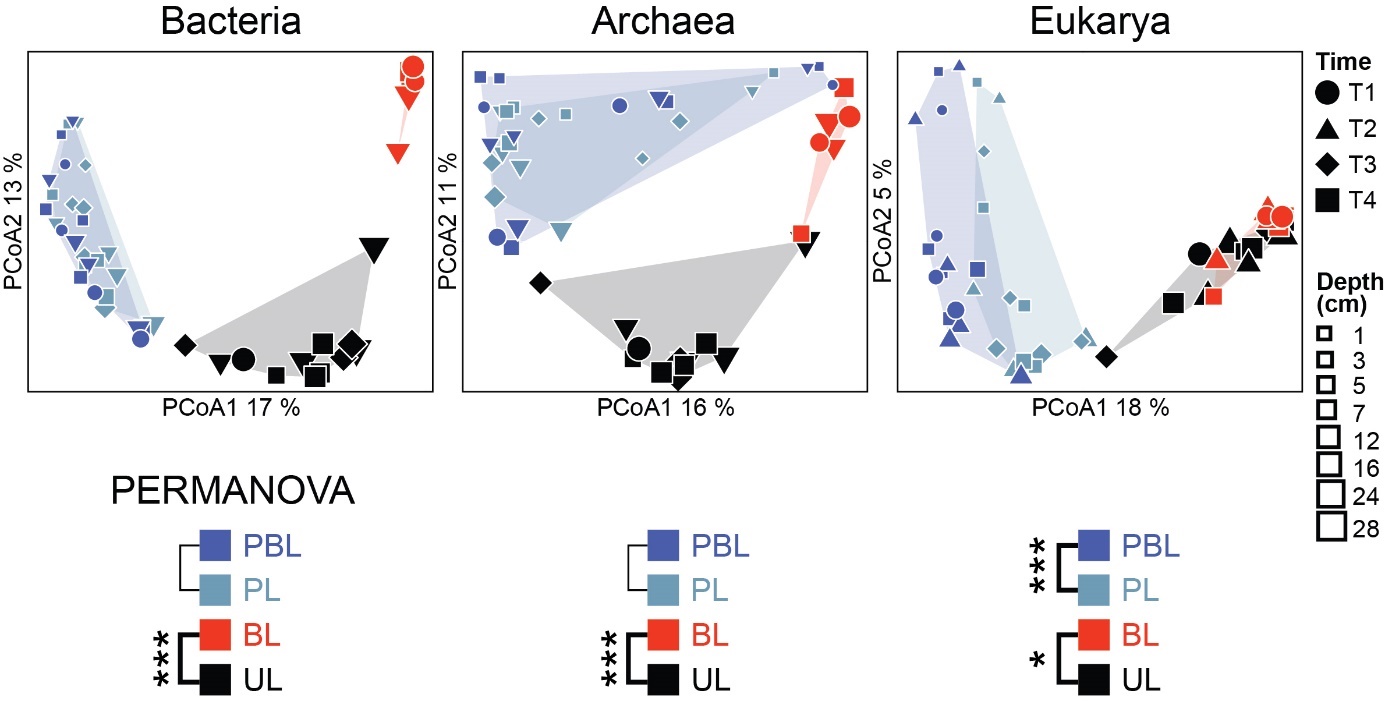

**Fig. S1.** Principal Coordinates Analyses (PCoA) based on unweighted Unifrac distance and PERMANOVA tests on bacterial, archaeal, and eukaryotic communities. In the calculation of community dissimilarity, unweighted Unifrac algorithm treats each taxon equally and is thus more sensitive at detecting differences in communities among low-abundance taxa.

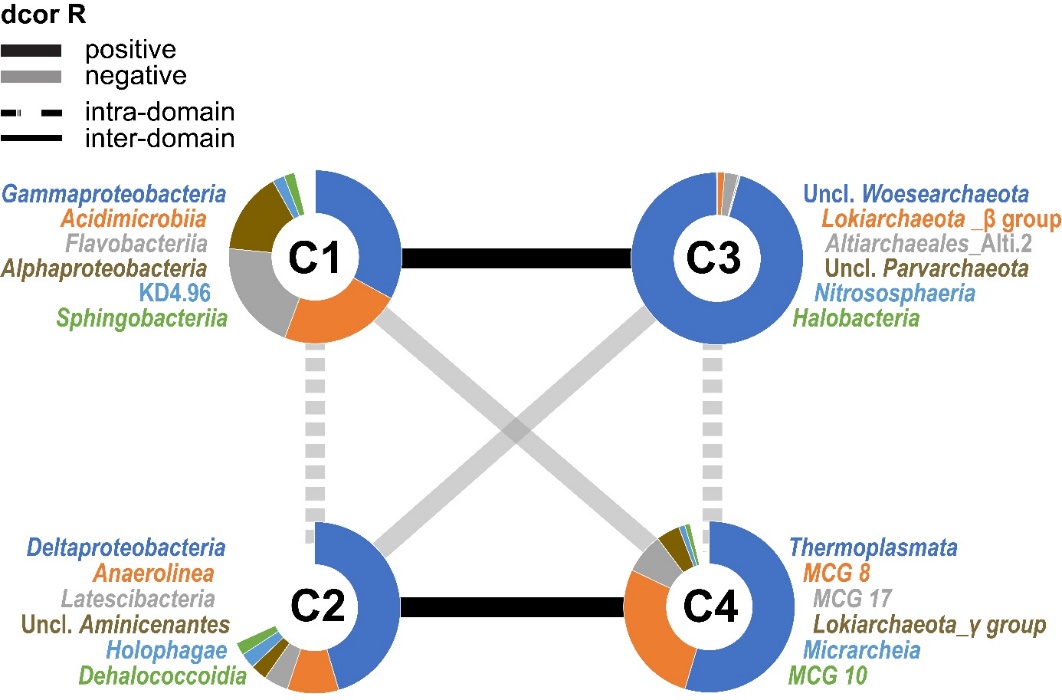

**Fig. S2.** Distance correlation test on clusters (C1-C4, see Fig. 5) in lugworm-free control sediments (999 bootstrap calculations). Only significant correlations (*p*<0.05) are reported.

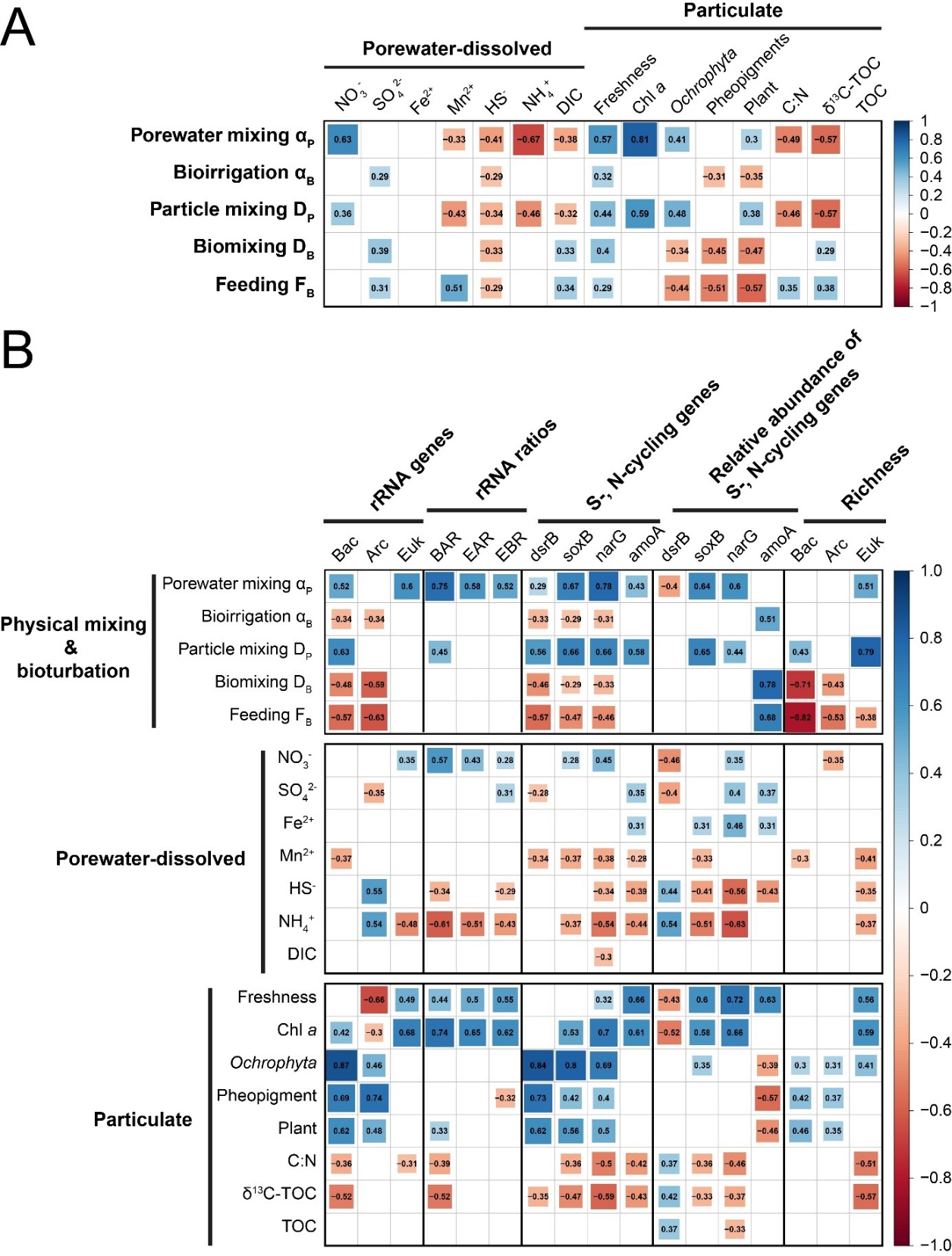

**Fig. S3.** Heatmap of relationships between (A) modeled mixing coefficients vs. geochemical data, and (B) modeled mixing coefficients and geochemical data vs. absolute and relative organismal gene abundances and ZOTU richness. All correlation values were calculated based on Pearson correlations, with r denoting the Pearson correlation coefficient, blue fields indicating significant positive correlations, and red fields indicating significant negative correlations. Blank fields indicate no significant correlation (p>0.05).

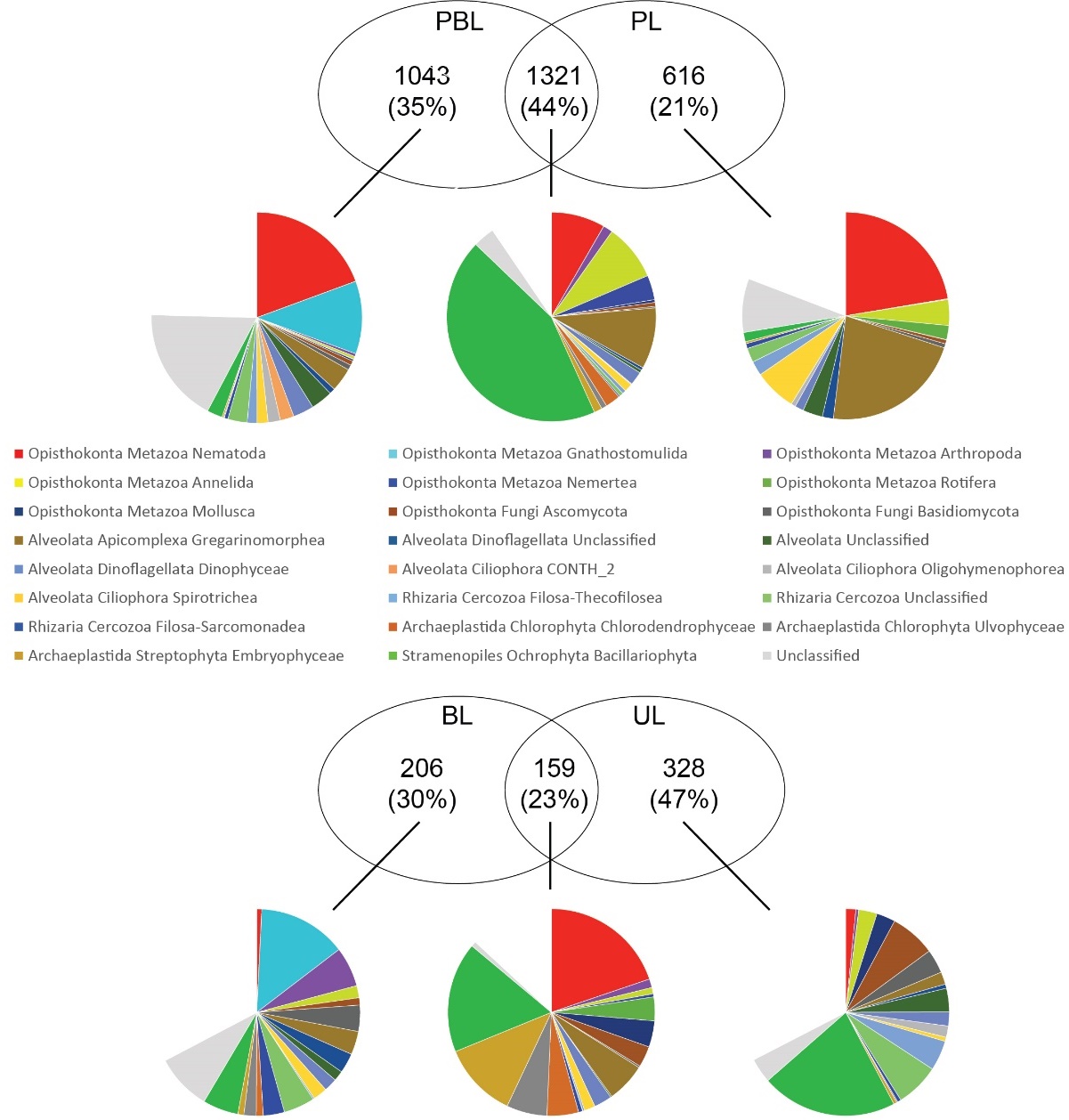

**Fig. S4.** Eukaryotic taxa that are unique to different sample types: physically and biologically impacted layers (PBL), physically impacted layers (PL), biologically impacted layers (BL), and undisturbed layers (UL).

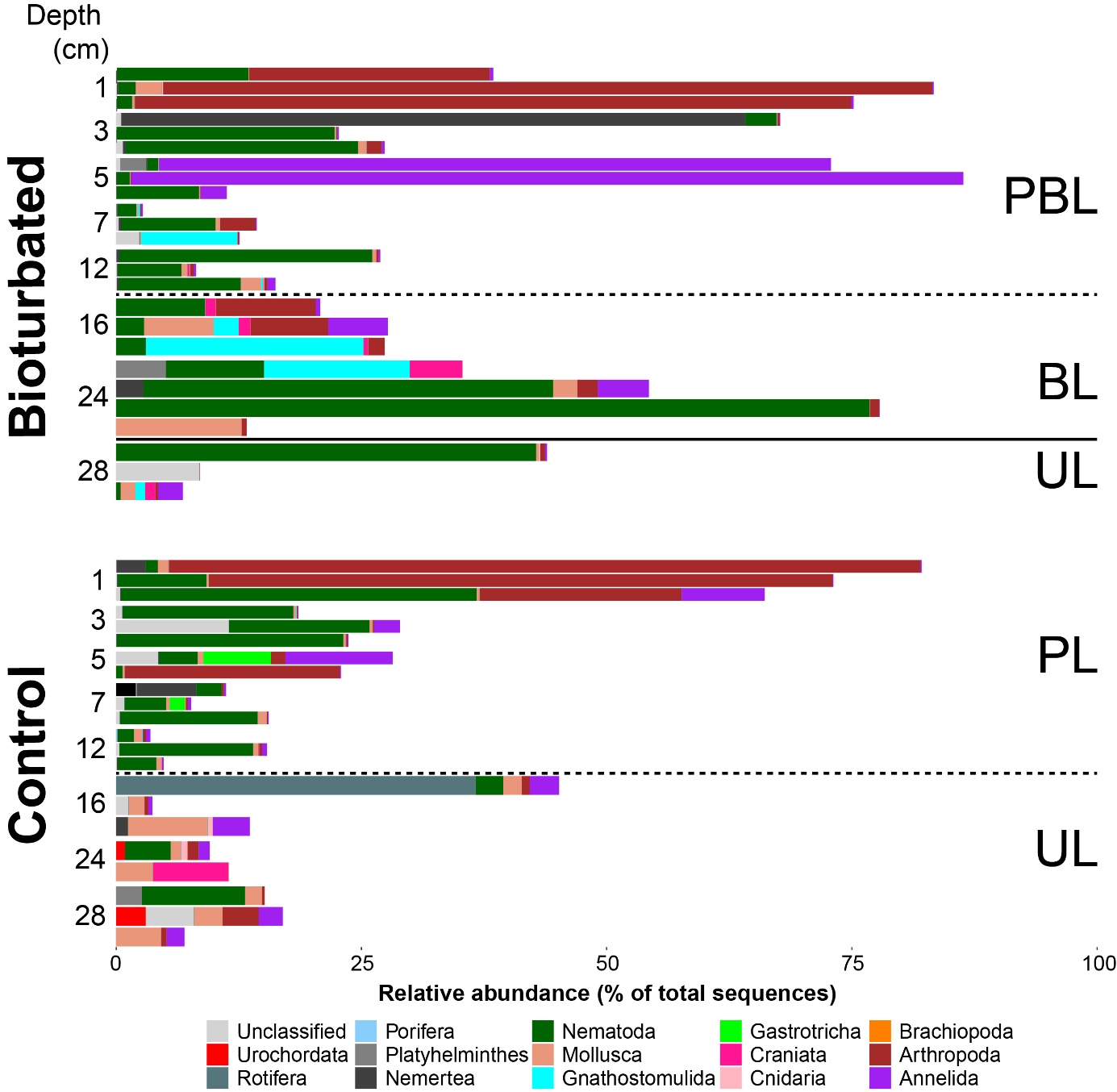

**Fig. S5.** Barcharts of relative abundances of Metazoan taxa versus sediment depth.

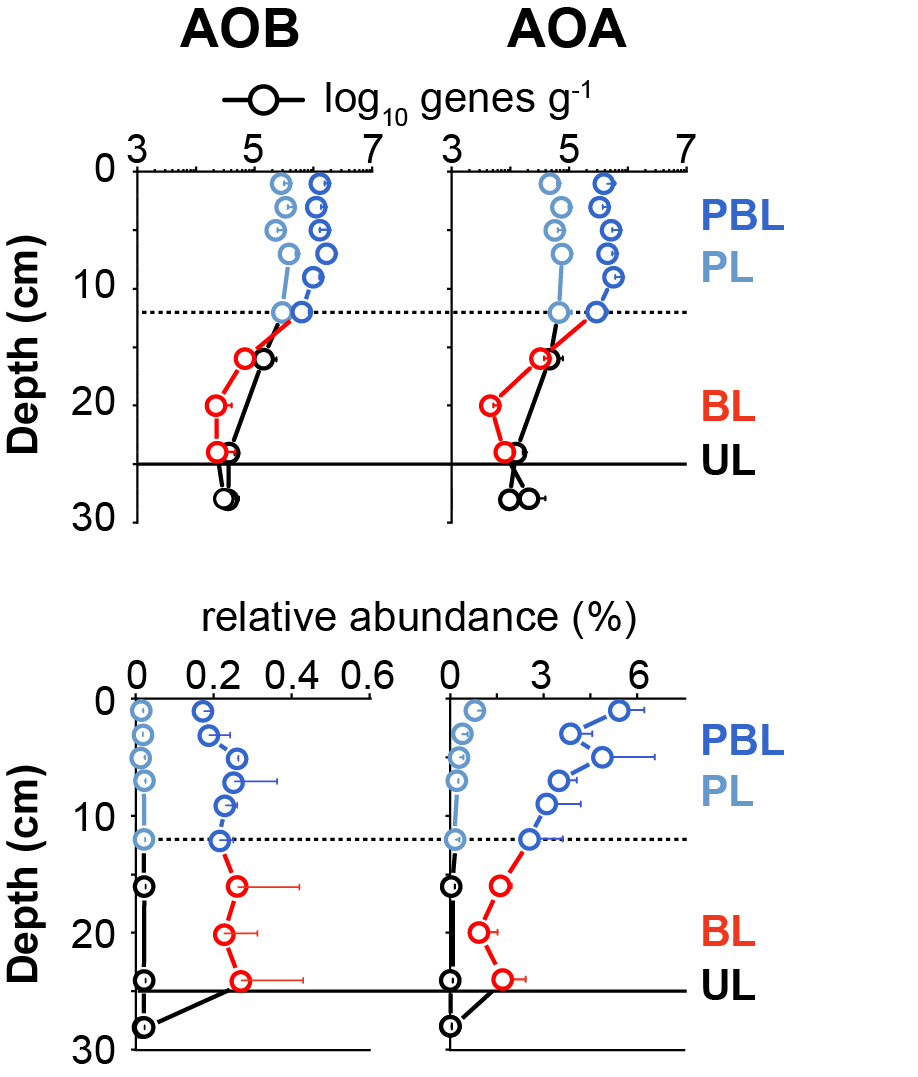

**Fig. S6.** Depth profiles of amoA gene copies that indicate the distributions of ammonium-oxidizing Bacteria (AOB) and ammonium-oxidizing Archaea (AOA) in sediment. Relative abundances of these genes in %, were calculated by dividing *amoA* gene copies by corresponding total 16S rRNA gene copy numbers. All values represent averages from four plots that were sampled at different time points (error bars denote standard deviations). [Abbreviations: PBL=physically and biologically impacted layer, PL=physically impacted layer, BL=biologically impacted layer, UL=undisturbed layer.]

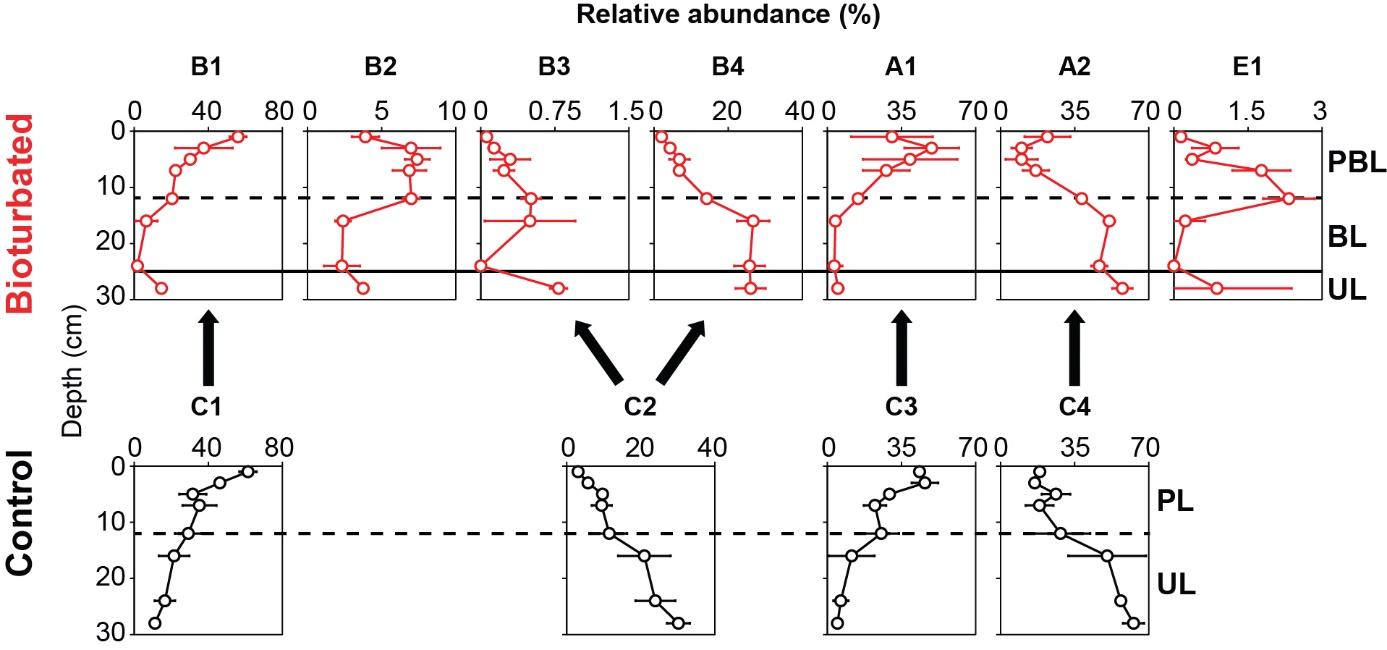

**Fig. S7.** Relative abundances of major network clusters plotted versus sediment depth.

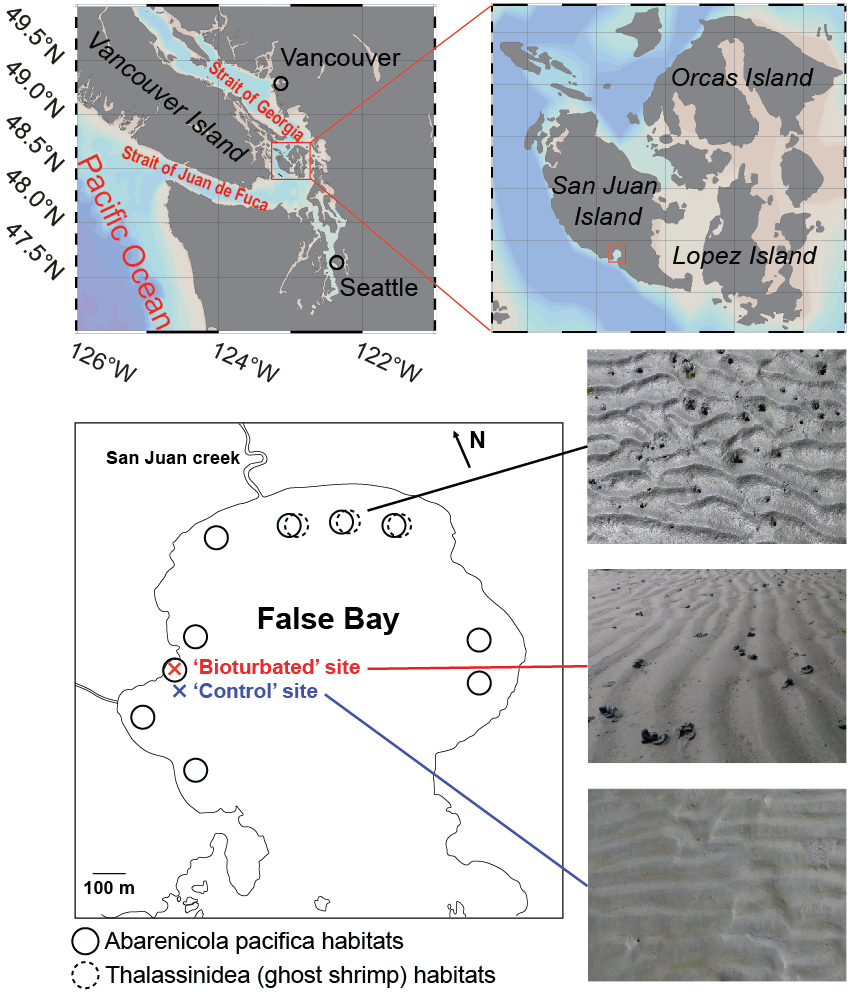

**Fig. S8.** Aerial maps and photos of sampling locations.

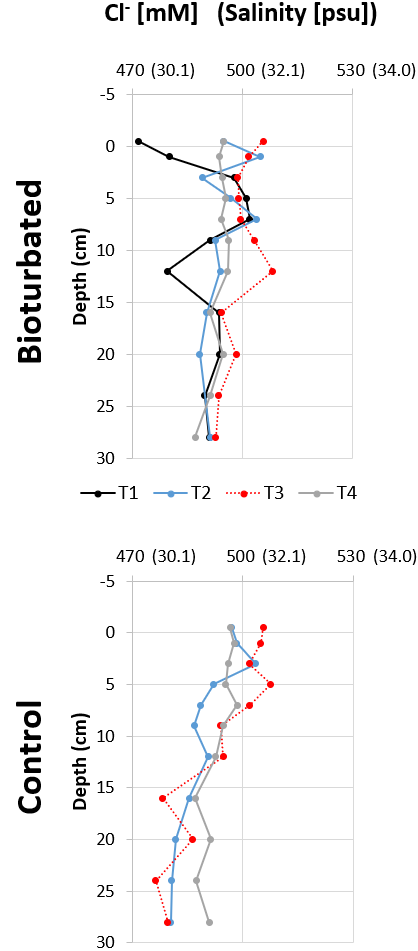

**Fig. S9.** Depth profiles of Cl^-^ concentration in bioturbated and control sediments. Values in parentheses are salinities calculated from Cl^-^ concentrations using the formula: Salinity (ppt) = Chlorinity (ppt) × 1.80655**.**

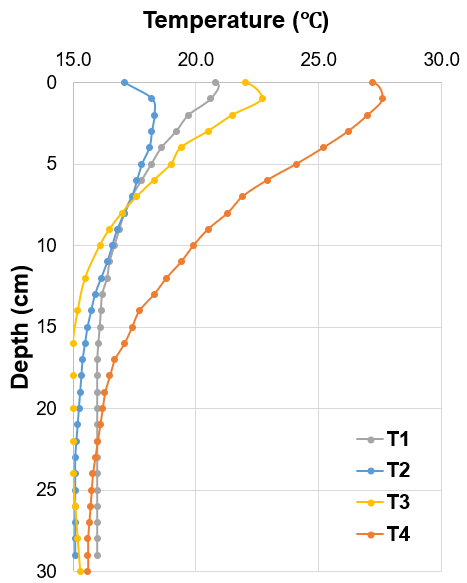

**Fig. S9.** Vertical temperature profiles in sediments before each sampling event (T1-T4).

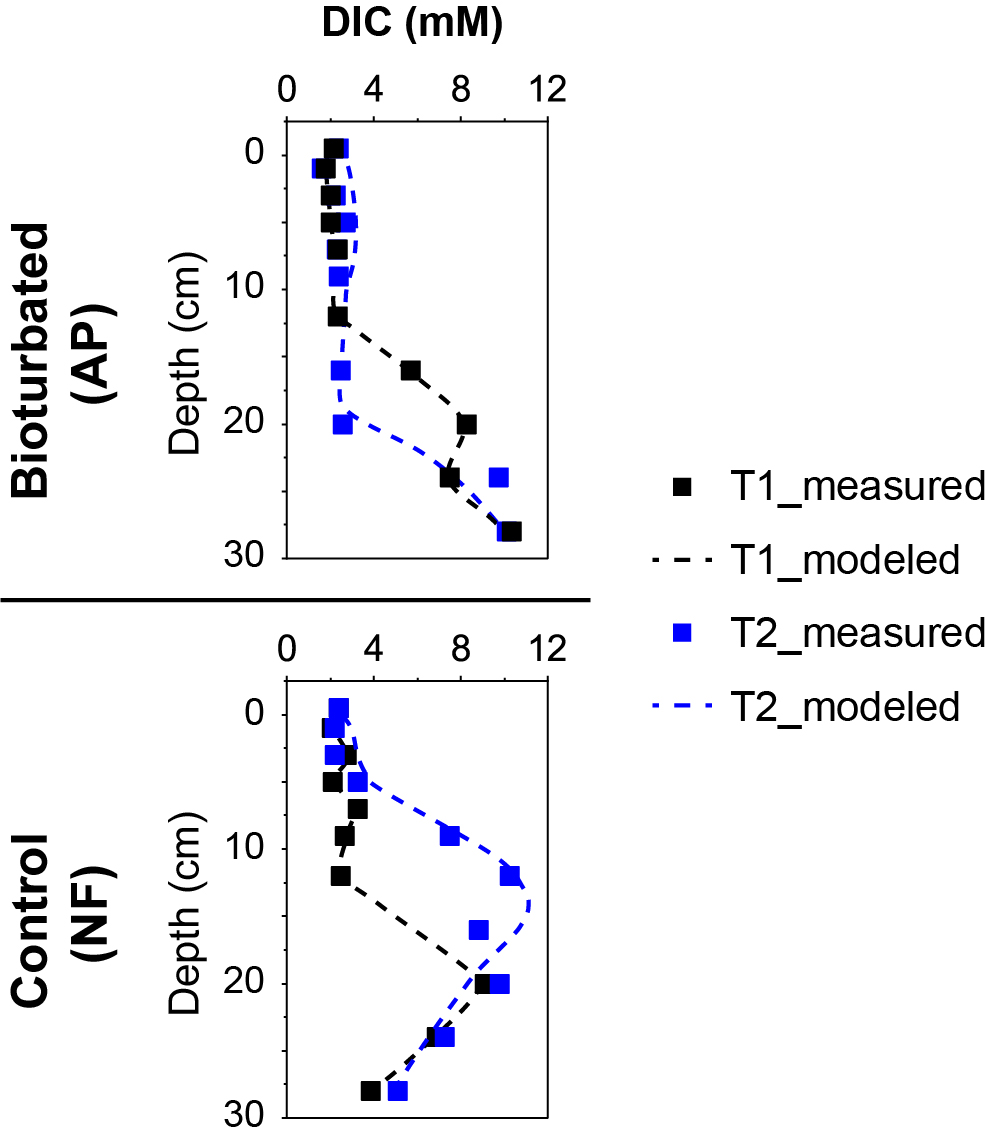

**Fig. S10.** Measured and modeled profiles of DIC in the refaunated (AP) and permanently defaunated treatments (NF). Modeled profiles were simulated using equation (S1) from “Modeled rates of porewater exchange by physical forcing and bioirrigation” in the Supplementary Text.

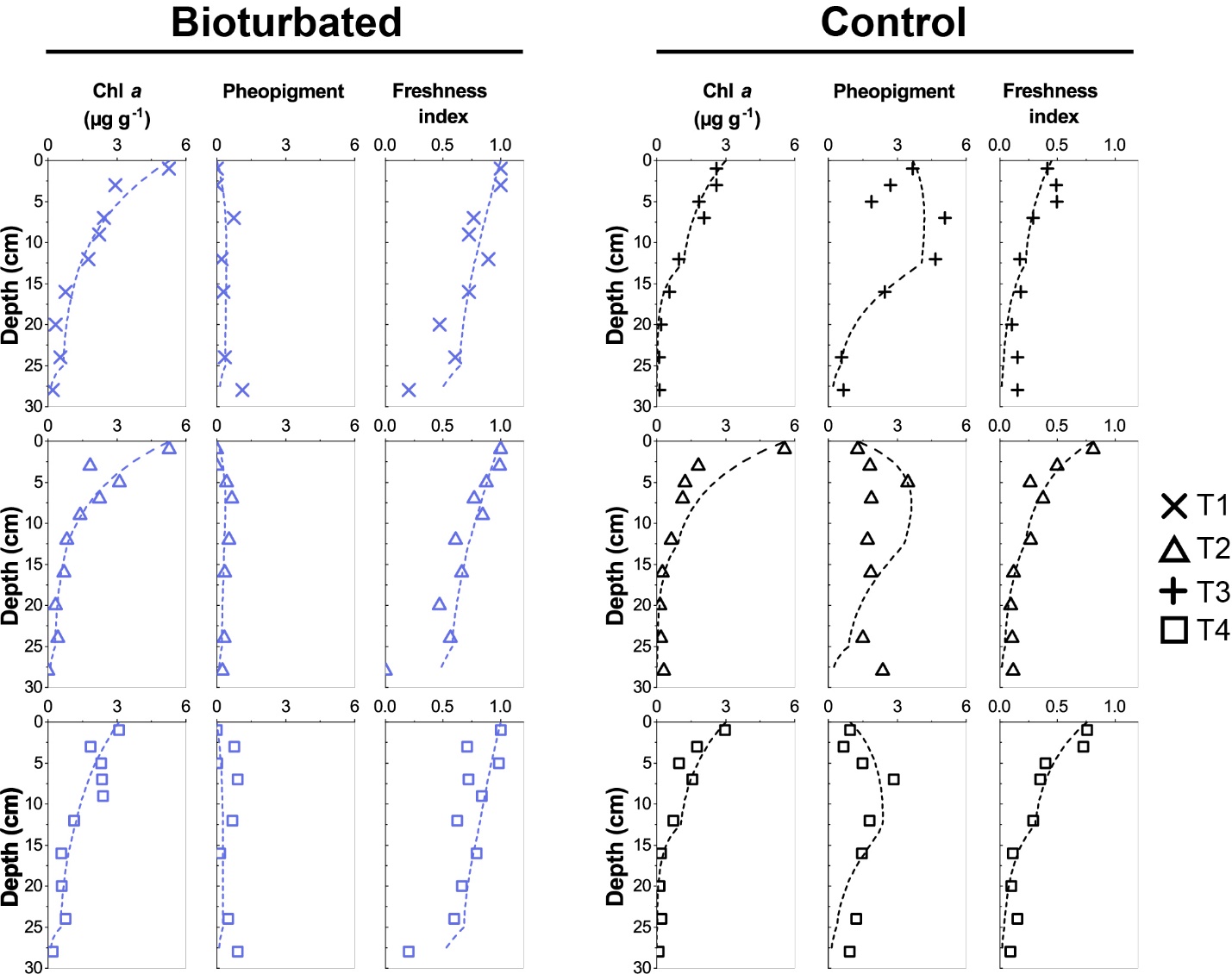

**Fig. S11.** Measured and modeled profiles of chl *a*, pheopigment, and freshness index in lugworm-inhabited (bioturbated) and lugworm-free (control) sediments. The modeled profiles were simulated using equations (2) and (3) from “Modeled rates of sediment mixing by physical and biological forcing” in the Supplementary Text.

Table S1. Overview of the primers and standards used for the qPCR and sequencing assays.

| **Target** | **Primer** | **Purposes** | **Sequence 5' - 3'** | **Target length (bp)** | **Annealing Temp.**  **(℃)** | **Reference** | **Pure cultures for standards** |
| --- | --- | --- | --- | --- | --- | --- | --- |
| **Archaeal** | Arc967F_mod | qPCR | AATTGGCGGGGGAGCAC | 144 | 55 | 10 | *Thermoplasma* |
| **16S rRNA** | Arc1059R |  | GCCATGCACCWCCTCT |  |  | 11 | *acidophilum* |
| **Bacterial** | Bac908F_mod | qPCR | AACTCAAAKGAATTGACGGG | 167 | 60 | 12, 13 | *Desolfotignum* |
| **16S rRNA** | Bac1075R |  | CACGAGCTGACGACARCC |  |  | 12, 13 | *phosphitoxidans* |
| **Eukarya** | All18SF_mod1 | qPCR & | TGCATGGCCGTTCTTAGT | 173 | 55 | 14 | *Tubifex tubifex* |
| **18S rRNA** | All18SR_mod1 | Sequencing | CTAAGGGCATCACAGACC |  |  | 14 |  |
| **Ochrophyta**  **rbcL** | Ochrophyta-F | qPCR | CGTTACGAATCTGGTGTAAT | 389 | 55 | 15 | *Stephanodiscus* sp. |
|  | Ochrophyta-R |  | GGAATACGCATATCTTCTAAACGTA |  |  | 15 |  |
| **Vascular** | rbcL h1aF | qPCR | GGC AGC ATT CCG AGT AAC TCC TC | 130 | 55 | 16 | *Salix* |
| **plant rbcL** | rbcL h2aR |  | CGT CCT TTG TAA CGA TCA AG |  |  | 16 | *chaenomeloides* |
| **narG** | NarG_1960F | qPCR | TAYGTSGGGCAGGARAAACTG | 90 | 60 | 17 | *Pseudomonas aeruginosa* |
|  | NarG_2050R |  | CGTAGAAGAAGCTGGTGCTGTT |  |  | 17 |  |
| **dsrB** | dsrB F1a-h | qPCR | CACACCCAGGGCTGG  CATACTCAGGGCTGG  CATACCCAGGGCTGG  CACACTCAAGGTTGG  CACACACAGGGATGG  CACACGCAGGGATGG  CACACGCAGGGGTGG  CATACGCAAGGTTGG | 362 | 56 | 18 | *Desulfobulbus propionicus* |
|  | dsrB 4RSI1a-f |  | CAGTTACCGCAGTACAT  CAGTTACCGCAGAACAT  CAGTTGCCGCAGTACAT  CAGTTTCCGCAGTACAT  CAGTTGCCGCAGAACAT  CAGTTTCCACAGAACAT |  |  | 18 |  |
| **soxB-1** | soxB-837Fa-i | qPCR | CAC AAC GGC ATG GAT GTN GA  CAC AAC GGC ATG GAC GTN GA  CAY AAC GGC TTC GAC GTS GA  CAY AAT GGC TTT GAC GTV GA  CAC AAC GGC TTT GAC GTV GA  CAT AAC GGC ATG GAT GTG GA  CAT GAT GGT TTT AGT GTT GA  CAT AAC GGC ATG CCG GTC GA  CAT GAT GGA TTC TCT GTG GA  CAC AAT GGT GCC GAT GTC GA  CAT AAC GGT ATG GAT GTT GA  CAT GAC GGG TTT GAC GTC GA | 333 | 60 | This study | *Thiobacillus denitrificans* |
|  | soxB-1170Ra-g |  | TT GAA RTT GCC SCG SCG RTA  TA GAA RGT ATC TCT TTT RTA  TA GAA ATT GTT GCG CCG RTA  TT AAA ATT ACC GCG TCG ATA  TC AAA ATT TCC CCG GCG ATA  TT AAA GTT GCC ACG ACG GTA  AA AAA TGT ATC ACG CTT ATA |  |  | This study |  |
| **amoA (AOA)** | Arch-amoAF | qPCR | STAATGGTCTGGCTTAGACG | 635 | 53 | 19 | *Nitrososphaera viennensis* |
|  | Arch-amoAR |  | GCGGCCATCCATCTGTATGT |  |  | 19 |  |
| **amoA (AOB)** | amoA-1F | qPCR | GGGGTTTCTACTGGTGGT | 491 | 55.4 | 20 | *Nitrosomonas europaea* |
|  | amoA-2R KS |  | CCCCTCKGSAAAGCCTTCTTC |  |  | 20 |  |
| **Archaeal** | ARC519F | Sequencing | CAGCMGCCGCGGTAAHACC | 396 | 63 | 21 | *Thermoplasma* |
| **16S rRNA** | ARC 915Rmod |  | GTGCTCCCCCGCCAATT |  |  | 10 | *acidophilum* |
| **Bacterial** | S-D-Bact-0341-b-S-17 | Sequencing | CCTACGGGNGGCWGCAG | 444 | 50-55 | 22 | *Desolfotignum* |
| **16S rRNA** | S-D-Bact-0785-a-A-21 |  | GACTACHVGGGTATCTAATCC |  |  | 22 | *phosphitoxidans* |

Table S2. Temperature protocols and corresponding time intervals for the qPCR assays.

| **qPCR step** | **Archaea 16S** | **Bacteria 16S** | **Eukarya 18S** | ***Ochrophyta rbc*L** | **Vascular plant rbcL** | ***nar*G** | ***dsr*B** | ***sox*B** | ***amo*A (AOA)** | ***amo*A**  **(AOB)** |
| --- | --- | --- | --- | --- | --- | --- | --- | --- | --- | --- |
|  | Time: min:ss  (Temperature: ℃) | | | | | | | | | |
| **1. Activation** | 05:00  (95) | 05:00  (95) | 05:00  (95) | 05:00  (95) | 05:00  (95) | 05:00  (95) | 05:00  (95) | 05:00  (95) | 10:00  (95) | 10:00  (95) |
| **2. Denaturation** | 00:10  (95) | 00:10  (95) | 00:10  (95) | 00:30  (95) | 00:30  (95) | 00:30  (95) | 00:30  (95) | 00:10  (95) | 00:30  (95) | 00:30  (95) |
| **3. Annealing** | 00:30  (55) | 00:30  (60) | 00:30  (55) | 00:40  (55) | 00:40  (55) | 00:30  (60) | 00:30  (56) | 00:30  (60) | 00:45  (53) | 01:00  (55.4) |
| **4. Polymerization** | 00:15  (72) | 00:15  (72) | 00:20  (72) | 00:30  (72) | 00:30  (72) | 00:15  (72) | 00:20  (72) | 00:15  (72) | 00:55  (72) | 00:55  (72) |
| **5. Acquisition** | 00:05  (80) | 00:05  (80) | 00:05  (80) | 00:05  (78) | 00:05  (80) | 00:05  (72) | 00:05  (82) | 00:05  (82) | 00:05  (72) | 00:05  (72) |
| **Repeat step 2-5: 50 cycles** | | | | | | | | | | |
| **6. Melting curve** | 01:00  (95) | | | | | | | | | |
|  | 1°C min^-1^  (60-95) | 1°C min^-1^  (60-95) | 1°C min^-1^  (55-95) | 1°C min^-1^  (55-95) | 1°C min^-1^  (55-95) | 1°C min^-1^  (65-95) | 1°C min^-1^  (55-95) | 1°C min^-1^  (55-95) | 1°C min^-1^  (55-95) | 1°C min^-1^  (55-95) |
